# Diet’s impact dictated by synonymous mitochondrial SNP interacting with nucleotype

**DOI:** 10.1101/2021.03.07.434274

**Authors:** Adam J. Dobson, Susanne Voigt, Luisa Kumpitsch, Lucas Langer, Emmely Voigt, Rita Ibrahim, Damian K. Dowling, Klaus Reinhardt

## Abstract

Epistasis between mitochondrial and nuclear genomes can modulate fitness effects of nutrition. Are these “mitonuclear” effects deterministic with regard to optimal nutrition? Which nutrients and genetic loci participate? Here, in fruitflies, we show that mitonuclear epistasis repeatably dictates fitness effects of dietary lipids and amino acids, with a tripartite interaction emerging as an unprecedented source of quantitative variation. We also observed diet-dependent developmental lethality and parental effects, but only in specific mitonucleotypes. Associating phenotype to mtDNA variation implicated a non-coding RNA (*mt:lrRNA*), suggesting a novel mechanism, with epistasis between mitochondria-derived regulatory factors and the nuclear genome producing qualitative differences in how diet impacts fitness.

**SUMMARY:** Combined action of nuclear genotype and regulatory loci on mtDNA produces major differences in fitness impacts of nutrition.

## Introduction

Nutrition and genotype are primary determinants of variation in health and biological fitness. They can also interact, resulting in different responses among genotypes to the same nutritional changes (*1, 2*). In humans, there is interest in leveraging this variation to optimize nutrition, by personalizing diet to individual consumers’ needs (*3*). To realize this ambition we must understand the genetic drivers of variation in response to nutrition, but this is challenging because independently segregating loci have non-additive, epistatic interactions (*4*), which may modulate response to nutrition, i.e. diet-by-genotype-by-genotype variation (*5*). Which genetic loci are involved? Mitochondria are critical metabolic hubs, with their own small genome, and variation in their function can contribute to variation in dietary optima. The mitochondrial genome segregates independently of the nuclear genome, and the combination of mitochondrial and nuclear variants can generate “mitonuclear” epistasis (*6, 7*). This epistasis is thought to occur because mtDNA is transcribed, processed and translated by nuclear-encoded proteins, and mtDNA-encoded proteins function in pathways and complexes that include nuclear-encoded proteins (*8*). Reciprocally, outputs of genetic variation in the nucleus depend on how mitochondrial metabolites and signals feed into broader cellular networks. Mitonuclear epistasis has been reported for numerous traits and processes (*9–11*), but diet-by-mito-by-nuclear (DMN) interactions are less well-characterized, despite evidence for mitochondrial modulation of nutrient signaling (*12*). So far, DMN interactions have been shown for development time, lifespan, fecundity and gene expression in *Drosophila melanogaster* (*5, 13–16*), but multiple important questions remain: (A) How much phenotypic variation do DMN interactions cause, relative to lower-order interactions (i.e. nuclear-diet, mitochondria-diet) and main effects (i.e. diet, nuclear, mitochondria)? (B) Is mitonuclear genotype deterministic for response to nutrition? In other words, when mitonuclear genotypes are replicated, are responses repeatable? (C) Parental nutrition can modulate offspring fitness, independent of offspring diet (*17*) - is this impact of nutritional variation subject to mitonuclear variation? (D) Which specific dietary nutrients cause DMN variation? Perhaps most importantly, (E) which mitochondrial polymorphisms underpin DMN interactions?

Here, in *Drosophila*, we study how variation among mitochondrial genotypes (mitotypes) modulates reproductive response to specific nutrients, in distinct populations of nuclear genotypes (nucleotypes), using a panel of fully-factorial mitonuclear pairings (mitonucleotypes). We study reproductive traits because of their relevance to biological fitness, expanding on preceding studies (*5, 13–16*) through multidimensional analysis of reproductive phenotype, and manipulating specific dietary nutrients (essential amino acids and lipid). We show that diets expected to promote fitness can in fact be lethal to specific mitonucleotypes. We also show that effects of parental nutrition on offspring performance are mitonucleotype-specific. This DMN variation was repeatable among independent genetic replicates. Effect sizes of DMN interactions were large for some traits, even exceeding those for diet:mitotype or diet:nucleotype interactions, implicating mitonuclear epistasis as a more important determinant of response to nutrition than variation in either genome alone, and showing that DMN variation can be a major source of phenotypic variation. Importantly, we observe DMN interactions among populations differentiated only by an mtDNA polymorphism was in a non-protein coding gene, long ribosomal RNA (*mt:lrRNA*), implicating a mitochondrial regulatory factor in DMN effects. Altogether these results suggest that mitonuclear epistasis can be a leading determinant of optimal diet, that this variation maps to putatively regulatory variants on mtDNA, and that the consequences can be a matter of life or death.

## Results

We produced *D. melanogaster* populations (Figure S1A) comprising replicated and fully-factorial combinations of mitochondrial and nuclear genomes from Australia, Benin and Canada (*A*, *B*, *C*, respectively). The crossing scheme was designed to produce distinct mitochondrial backgrounds bearing equivalent pools of standing nuclear variation, by introgressing populations either reciprocally or to themselves. For brevity, we abbreviate population names, giving mitochondrial and then nuclear origin (e.g. *AB* = Australian mitochondria, Beninese nuclei). Each combination was triplicated (e.g. *AB_1_, AB_2_*, *AB_3_*) at the beginning of the introgression and maintained in parallel for more than 160 introgressions, altogether generating 27 populations, comprising nine triplicated mitonucleotypes. We confirmed that this design (A) replaced original nucleotypes with donor nucleotypes (Figure S1B, Figure S1C), at frequencies from 94.4% to 99.9% (Figure S1F, Supplementary Text), and that (B) nucleotype replacement was equivalent in the presence of distinct mitotypes (Figure S1B, Supplementary Text). These analyses confirmed that the panel of populations was suitable for studying the behaviors of distinct mitotypes in populations with comparable nuclear genomic diversity. These analyses also indicated that the majority of mitotype diversity was represented by populations bearing only Australian or Beninese mitochondria (Figure S1B, Supplementary Text), and so populations with Canadian ancestry were excluded from further study, to reduce the total number of populations for phenotyping from 27 to 12 (Supplementary Text). We then examined the fitness response of these populations (*AA_1-3_, AB_1-3_, BA_1-3_ and BB_1-3_*) to dietary manipulations, applying both an established manipulation that promotes fecundity by enriching essential amino acids (EAA), and a novel manipulation that represses fecundity by enriching plant-based lipids (Figure S2A, Supplementary Text). Feeding these EAA-enriched and lipid-enriched diets to the focal panel of 12 populations (Figure S2C, Supplementary Text) revealed mitonuclear variation in fecundity response (Figure S2D, Supplementary Text), which could not be accounted for by caloric density (Figure S3B, Supplementary Text), revealing for the first time specific nutrients that are sufficient to elicit DMN variation (Figure S3B, Supplementary Text). This fecundity variation was both (A) technically repeatable among replicate experiments (Figure S3A, Supplementary Text), suggesting that introgression maintained equivalent diversity across generations; and (B) biologically repeatable among replicate lines (Figure S2E, Supplementary Text), suggesting that the repeatable genetic variation in the lines led to predictable and repeatable phenotypic variation.

### Large effects of DMN interactions on multiple fitness parameters

Encouraged by initial fecundity results, we characterized a more extensive panel of fitness traits. In a total of >25,000 individual flies we assayed fecundity, fertility and development time - traits that have previously been shown to be sensitive to DMN variation (*13, 16*) - as well as number of adult progeny as a direct fitness measure. We varied feeding on experimental diets in two different ways (Figure 1A). We hypothesized that there might be mitonucleotype-dependent variation in response to (A) chronic dietary change for both parents and offspring, and (B) parental diet, which can influence offspring health independent of offspring diet (*17, 18*). Therefore, eggs were laid and developed either in a chronic feeding paradigm, in which both parents and offspring were fed experimental diets; or a parental feeding paradigm, in which diets were fed to parents before eggs were laid and developed on a standardized medium, distinct from parental diet. In the latter context, DMN interactions can only result from a parental effect. To ensure genetic consistency, the same parents were used in each paradigm, by laying eggs for 24h after one week on experimental media (chronic paradigm), then switching to a universal standardized medium for another 24h of egg laying (parental paradigm). The parents developed on a standardized medium that was distinct from all experimental media, to ensure that any novelty effects were distributed evenly among experimental conditions.

**Figure 1.**
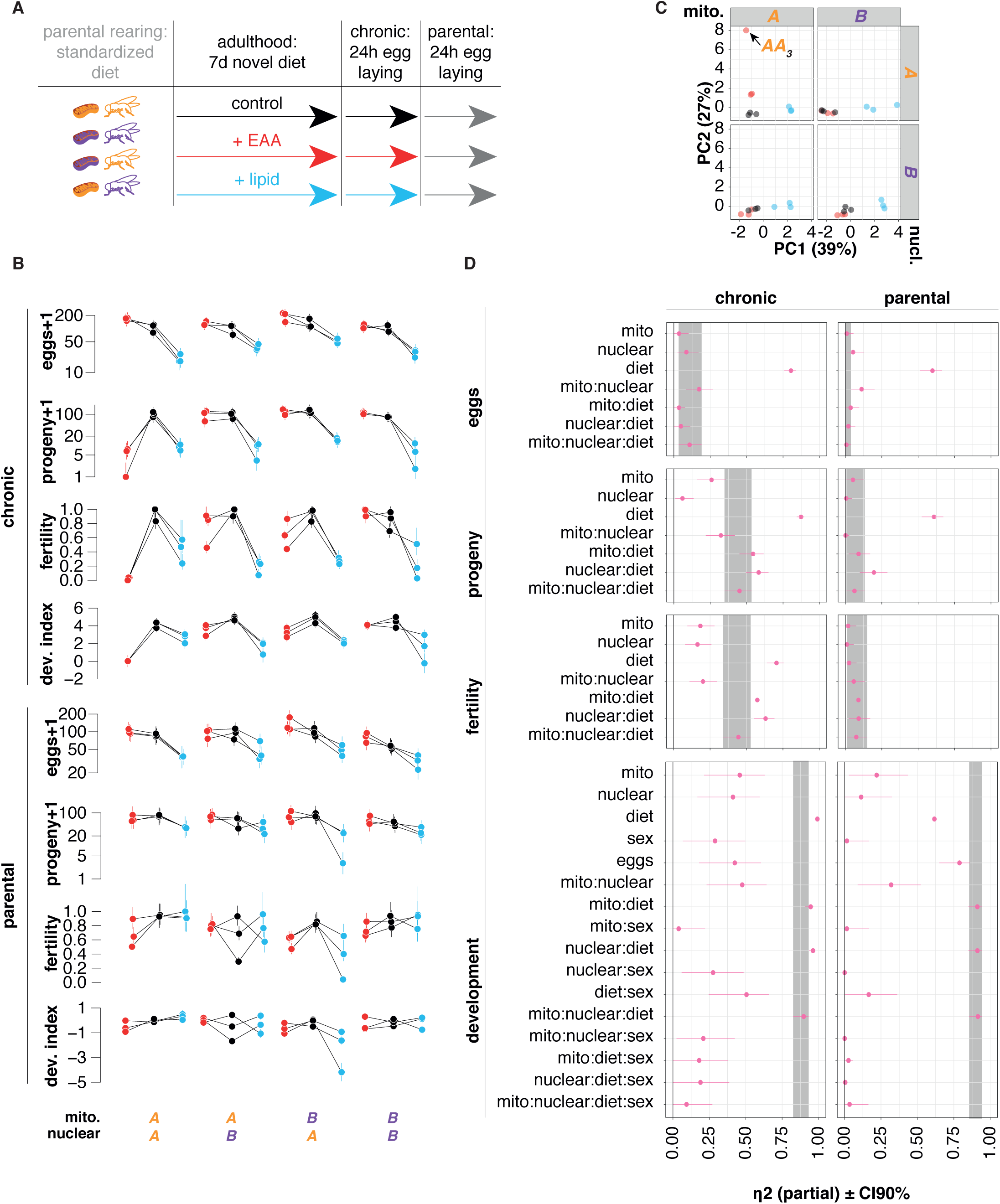
Large and repeatable impacts of mitonucleotype on response to chronic and parental nutritional change. **A.** Key and experimental design. Flies were reared from egg to adult on rearing food, and allocated at random to experimental media 6-48h after eclosion, at a density of five of each sex per vial. After seven days, flies laid eggs on fresh food for 24 hours, followed by a further 24h son standardized rearing medium. **B.** Repeatable mitonuclear variation in response to chronic and parental changes in nutrition. Panels show EMMs for trait indicated on y-axis. Feeding paradigm and mitonuclear variation are indicated at the top of the plot. Colors encode diet as per key. Development index shows EMMs for Cox mixed-effects models of proportion eclosed over time, excluding sex from plot. Development data are plotted in full as Kaplan-Meier plots in Figure S4. Note the exclusion of EMMs for development of genotype *AA_3_* in chronic feeding: EAA lethality prevented meaningful estimation. **C.** PCA shows ordination of populations according to mitotype, nucleotype, and diet. Results shown from one PCA of scaled and mean-centered EMMs, split by facets per genotype, with mitotype split by rows and nucleotype split by columns. **D.** Effect sizes for mitonuclear modification of responses to nutrition. Panels show effect size estimates (partial η^2^) for traits indicated on left, in feeding paradigms indicated at top. Partial η^2^ calculated from GLMMs (fecundity, progeny, fertility) or Cox Mixed Models (development). Grey bands indicate confidence interval (90%) for diet-mito-nuclear terms. For all traits in the chronic feeding paradigm, diet consistently had the largest effect size, but diet-mito-nuclear effect size was either greater than or equal to mito-diet and nuclear-diet terms, and also larger than mito or nuclear main effects. In the parental feeding paradigm, for fertility and development but not fecundity or progeny, diet-mito-nuclear effect size was either greater than or equal to other genetic modifiers of response to nutrition, and by ranking greater than main mito or nuclear effects.

To visualize patterns of DMN variation, and repeatability among replicate mitonucleotypes, we calculated Estimated Marginal Means (EMMs (*19*), Figure 1B). DMN variation was visually apparent for all traits (Figure 1B – noting exclusion of genotype *AA_3_* from development in chronic paradigm, see Supplementary Text). Statistical models revealed DMN interactions for all traits in each feeding paradigm (GLMMs for fecundity, progeny and fertility; Cox mixed-effects models for development time), except for fecundity in the parental feeding paradigm (Table S5). No trait varied as a linear function of caloric density (Figure S5), and so impact of diet was modelled as an unordered factor. For development time models, we also included interactions with offspring sex, because of reports of sex-biased mito:nuclear variation (*16*). However, sex did not modify genotype-by-diet interactions (all sex interactions with diet, mitochondria, or nuclear background: p>0.05).

Chronic lipid feeding was deleterious for all traits, but mitonuclear genotype determined magnitude; chronic EAA feeding promoted fecundity but, surprisingly, reduced fertility (Figure 1B). Consequently, progeny counts on EAA did not exceed counts on control diet, suggesting that fitness was not enhanced by EAAs. The magnitude of the fecundity benefits and fertility costs were again mitonucleotype-dependent. In *AA* populations, uniquely, progeny count after chronic EAA feeding was even lower than after chronic lipid enrichment, to near lethality in *AA_3_* (Figure 1B), but this sensitivity was rescued by replacing either the mitochondria (*BA*) or nuclei (*AB*) (Figure 1B), confirming a mitonucleotype-specific effect. Mitonuclear incompatibility is widely reported (*20*), but our data indicate that it can be diet-dependent, under nutrient-enriched conditions that we had expected to promote fitness.

DMN variation was also apparent in the parental feeding paradigm, albeit less pronounced than after chronic feeding. Lipid was less universally toxic. *AA* flies even exhibited a benefit of parental lipid feeding, developing on average one day earlier (Figure S4), but again this was not evident in *BA* or *AB* flies (Figure 1B), confirming another mitonucleotype-specific effect.

To estimate repeatability for all traits, we used the same approach as in our initial fecundity experiments (Supplementary Text), and found among-replicate repeatability both within each diet and in response to nutrient enrichment, for all traits (Figure S6).

To distill the complex patterns of variation in this multi-dimensional dataset, we performed PCA on EMMs for all traits on all diets in all populations, for an integrative view of how mitonucleotype determined response to diet (Figure 1C). Control or EAA-fed flies separated from lipid-fed flies on the first principal component. However the second principal component revealed an orthogonal response to EAAs in *AA* mitonucleotypes, particularly in line *AA_3_* (Figure 1C). This analysis was consistent with an interpretation that DMN effects induce coordinate changes in multiple traits, leading to an altered phenotype overall.

We used these data to address one of our main questions: Is mitonuclear epistasis deterministic for the impact of nutritional variation? We calculated a measure of effect size (partial η^2^) for DMN interactions, and compared to lower-order effects (Figure 1D), anticipating that diet would be the largest source of variation for most traits, but that this this might be modified by mitonucleotype (i.e. large DMN terms). However we were surprised to find that the magnitude of DMN effects approached or equaled diet’s effect for some traits, indicating a major source of variation. In the chronic feeding paradigm, DMN effect sizes were greater than or equal to mito:diet and nuclear:diet effects for egg laying, progeny and fertility (Figure 1D). For development in both paradigms, DMN effect sizes were large, on par with diet, diet:mitotype, and diet:nucleotype; revealing DMN interactions as a major source of variation for developmental impacts of chronic dietary change in these flies. Fertility effects were more pronounced after chronic feeding than after parental feeding, but in both paradigms DMN effect sizes were ∼75% of diet’s main effect, as were mito:diet and nuclear:diet terms, suggesting that these factors do not simply modulate effect of diet, but their interaction with diet is a substantial source of variation outright. In fact, in the parental nutrition paradigm, DMN effect sizes for fertility outranked the main effect of diet, although with overlapping confidence intervals and modest effect size. However, for development, effect sizes for DMN terms exceeded nearly all other terms (exception for the lower-order diet:mito and diet:nuclear interactions), without overlapping confidence intervals (Figure 1D). The magnitude of effect on this trait exceeded even the main effect of diet, suggesting that dietary regulation of this trait could not be properly understood without accounting for mitonucleotype. We also validated our effect size calculations orthogonally, by assessing how a range of alternative models described the data, and by calculating variance explained (r^2^) by each model, which gave congruent results (Supplementary Text). These analyses suggest that mitonucleotype can modulate response to dietary variation, and the emergent interaction can produce as much phenotypic variation as the main effect of diet.

### Associating DMN interactions with mitochondrial polymorphisms implicates regulatory variation

The second major question of our study was: what are the putative causes of DMN interactions? Mitochondrial polymorphisms underlying DMN interactions are largely uncharacterized. We examined our populations’ mitochondrial SNPs in detail, to ask if phenotypic variance segregated according to mitonucleotype. mtDNA SNP frequencies were stable over time (Supplementary Text, Table S8), suggesting we could use these flies in the same way as established genetic mapping populations (*21*), because introgression maintained variation and therefore phenotypic variation could be associated with preceding sequence data. We found 27 SNPs, in both protein coding and non-coding regions of mtDNA (Table S3), but only 2/27 SNPs were expected to change amino acid sequence (Table S3), suggesting that synonymous and regulatory polymorphisms might underlie DMN interactions (Supplementary Text). To reveal how major mtDNA alleles co-segregated, and how they intersected with nucleotypes, we performed hierarchical clustering of major alleles, and examined how the clustering grouped populations (Figure 2A). This separated *A* mitotypes from *B*, validating our initial approach of encoding mitotype by geographic origin (Figures 2, 3). However, nested within this higher-level grouping, more granular among-line differentiation was evident, with five distinct clusters of unique mitotypes. These mitotypes were not nested within nucleotype, and some co-occurred with both *A* and *B* nucleotypes. We confirmed this pattern of co-occurrence orthogonally, with network analysis (Supplementary Text). These patterns of co-occurrence suggested that differential responses to diet could be associated to the distinct mitotypes. To generate a final, sequence-informed mitonucleotype assignment, we concatenated sequenced-based mitotype with nucleotype (i.e. A or B, since our sequencing data suggested nuclear homogenization independent of mitotype (Figure S1)). This revealed eight distinct mitonucleotypes. We re-analyzed our phenotype data with respect to these new mitonucleotype assignments, fitting models that tested for interaction of mitonucleotype and diet, which confirmed distinct responses to diet (Supplementary Text, Figure S8).

**Figure 2.**
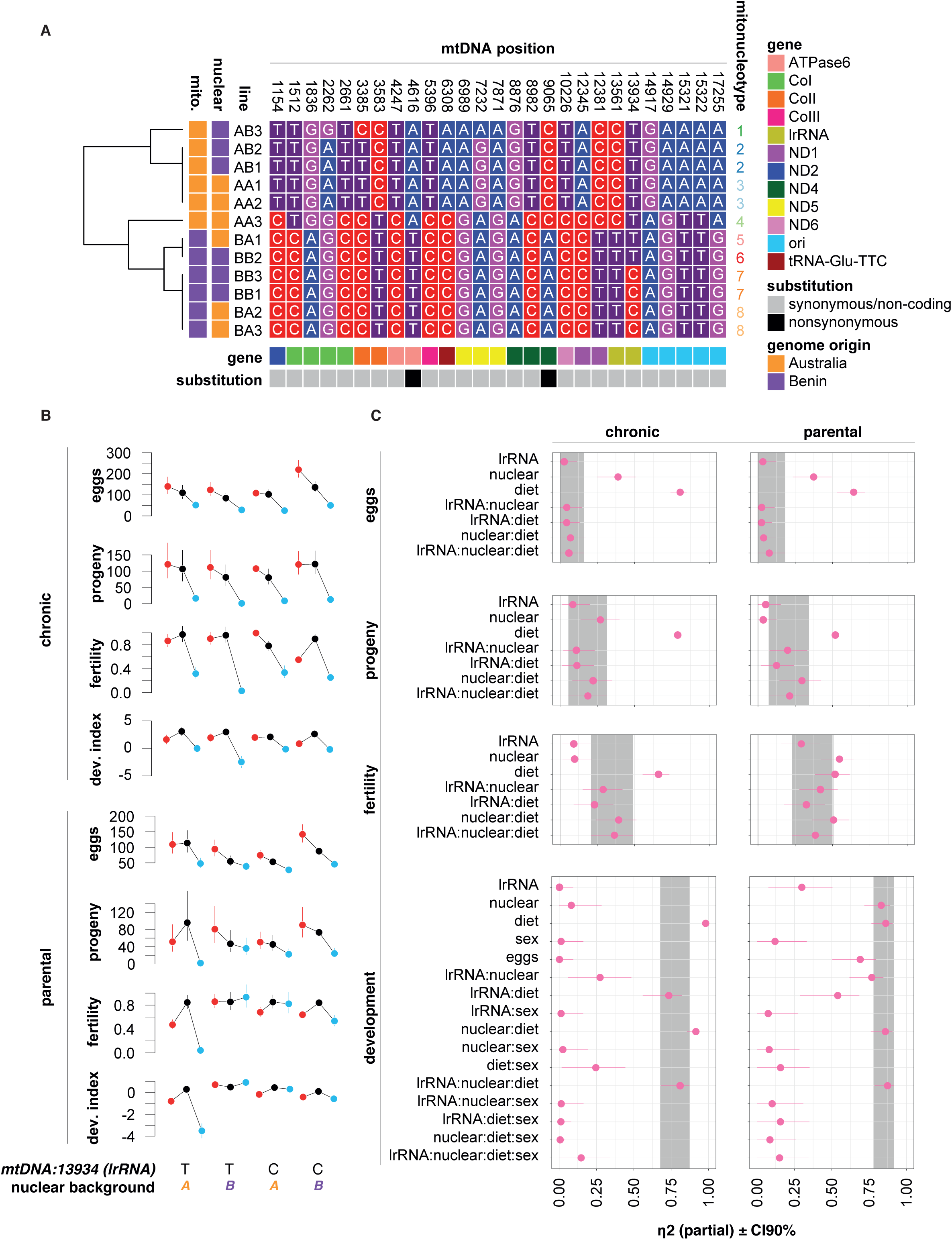
A C/T polymorphism in mitochondrial *long ribosomal RNA* substantially alters nucleotype-specific response to dietary variation. **A.** Segregation of major alleles for significantly differentiated mitochondrial SNPs. Heatmap shows nucleotide identity at positions in mitochondrial genome indicated at top. Gene for each position and SNP classification (synonymous/non-synonymous) indicated by color bar at top, and geographic origin of mitochondrial and nuclear genomes indicated on right. Hierarchical clustering (dendrogram on left) shows separation of SNPs by geographic origin, with five constituent clusters. Concatenating SNP clusters with nucleotype reveals eight mitonucleotypes, indicated to right. **B.** Trait EMMs among subset of populations with mitochondria varying only by a C/T substitution in *mt:lrRNA* (position 13934). Significant SNP-by-nucleotype-by-diet effects were observed for progeny, fertility and fecundity (Table S10). Panels show EMMs for traits and feeding conditions indicated on left, with mitonucleotype indicated at bottom. EMMs calculated from GLMMs (fecundity, progeny, fertility) or Cox mixed models (development). Equivalent plot for all populations in the study shown in Figure S8. **C.** Effect sizes for interaction of mt::lrRNA (13934) C/T polymorphism, nucleotype and diet. Panels show effect size estimates (partial η^2^) for traits indicated on left, in the two feeding paradigms indicated at the top. η^2^ calculated from GLMMs (fecundity, progeny, fertility) or Cox Mixed Models (development). Grey bands indicate confidence interval (90%) for diet-mito-nuclear terms. Effect sizes suggest that, for progeny and fertility of these populations (mitonucleotypes 5-8), the variation resulting from the interaction of lrRNA polymorphism, nucleotype and diet is equivalent to standing genetic variation from nucleotype and lrRNA polymorphism; and also an equivalent determinant of response to diet.

We were particularly interested by the paucity of nonsynonymous mtDNA polymorphisms, which suggested roles for putatively regulatory variants. We noticed that a subset of populations (mitonucleotypes 5, 6, 7, and 8) (Figure 2A), were distinguished by only one mitochondrial polymorphism. This was a C/T polymorphism in *mt:lrRNA* (mtDNA position 13934), which occurred at high frequencies (between 0.99 and 1, Table S3) in each nuclear background. The presence of only one mtDNA SNP in these lines (contrasting other populations, which bore confounding variation at other positions) suggested that any mitonuclear or DMN variation in this subset of lines was attributable to the *mt:lrRNA* SNP. *mt:lrRNA* is not known to encode protein, so this SNP could provide an illustrative example of the potential for mitochondrial regulatory variation to drive DMN variation in phenotype.

Plotting phenotype data from mitonucleotypes 5, 6, 7 and 8 (Figure 2B) revealed qualitative variation in fertility. In populations with the *mt:lrRNA* T allele, chronic EAA feeding decreased fertility, in both nucleotypes (mitonucleotypes 5 & 6). However, the *mt:lrRNA* C allele unleashed nucleotype-dependent responses to diet: C allele populations with nucleotype A showed decreased fertility after EAA feeding (mitonucleotype 8), but increased fertility with nucleotype *B* (mitonucleotype 7) (Figure 2B). Indeed, mitonucleotype 7 was the only mitonucleotype that increased fertility upon EAA feeding.

In the parental feeding paradigm, nucleotypes *A* and *B* responded to diet equivalently in the presence of the *mt:lrRNA C* allele (mitonucleotypes 7 & 8). However, nucleotype-specific responses to diet were unleashed by the *mt:lrRNA* T allele: fertility was impaired by parental feeding on either EAA or lipid in the presence of nucleotype *A* (mitonucleotype 5), but not in the presence of nucleotype *B* (mitonucleotype 6). This altered fertility had apparent consequences for progeny count and development index (Figure 2C). Statistical tests (Table S10) confirmed interactions of the *mt:lrRNA* polymorphism, nucleotype, and diet, for all traits except egg laying. This exclusively post-embryonic variation indicated impacts on offspring performance, but not parental reproductive effort. Effect size calculations (Figure 2C) suggested that the lrRNA:nucleotype:diet interaction was an important source of variation among these populations. For fertility during chronic feeding, lrRNA:nucleotype:diet effects were bigger even than for nucleotype. Most strikingly, for development in both feeding paradigms, effect size for the lrRNA:nucleotype:diet interaction was large, approaching or even equal to main effects of diet. Altogether these results suggest that epistasis between nucleotype and a SNP in non-coding mtDNA can dictate response to diet, which can produce more phenotypic variation than the main effects of mitochondrial or nuclear genotype, and can equal the effect of diet.

## Discussion

Predicting phenotype from genotype is a long-standing challenge. To this end, genome-wide association studies (GWAS) have flourished. Two over-arching findings of the era of GWAS are that non-coding variation is more important than previously expected; and that additive effects of independently-segregating variants do not fully explain quantitative trait variation (*4, 22*). This latter finding implies “missing heritability”, suggesting that additional processes are at work. Two hypothetical explanations are that genotype-by-genotype epistasis (G*G) and genotype-by-environment (G*E) interactions create non-additive effects. Speculation about epistasis has led to the “omnigenic model”, which posits that variation in a given trait is likely explained by G*G between a few “core genes”, and the sum effect of many (or all) small-effect variants throughout the rest of the genome (*4, 22*). Mitonuclear interactions may be a useful illustration of the omnigenic model, with epistasis between the few genes on the mitochondrial genome and the sum of nuclear genomic variants producing substantial phenotypic variation (*7*). We are not aware of any prior attempt to explicitly associate specific mitochondrial SNPs with these interactions, and our results suggest that non-coding variants on mtDNA can play a role. It remains to be seen if non-coding mtDNA variation is as important as non-coding nDNA variation appears to be, though our effect size calculations (Figure 2C) indicate potentially large roles. More generally, we have added to the growing body of evidence for diet-mito-nuclear interactions (*5, 13, 16*), showing that outcomes of mitonuclear epistasis are environment-dependent. Thus, altogether, DMN interactions show many of the hallmarks of a major source of phenotypic variation, and we have demonstrated this for *Drosophila* fitness traits. The Darwinian view that reproduction subjugates all other processes, and the central role of mitochondria in cellular function, suggest that these interactions may be important but underappreciated sources of variation for many further traits, and not just in flies.

While we partitioned phenotypic variance to SNPs, we have not attempted systematic GWAS. In general, GWAS to test genome-wide epistasis are not tractable, due to the enormous sample that would be required to maintain statistical power. For mitonuclear epistasis, testing consequences of interactions between “only” every mtDNA variant and every nuclear variant would be simpler, but an enormous sample would still be required. However, if the omnigenic hypothesis is correct (*4*), such a systematic approach would fail to recognize the underlying biology, which is better modelled as epistasis between a subset of core genes (i.e. mitochondrial genes) and nuclear genomic background (e.g. represented by “background” as in our study; or alternatively dimension reduction, pedigrees, marker loci, or pathway-level variation). If candidate mtDNA variants can be identified, methods to test their role conclusively, by mtDNA editing, are on the horizon. mtDNA editing is in its infancy (*23*), but would facilitate powerful tests of how mitochondria affect outputs of nuclear variation, including response to diet. An additional question raised by our study is the mechanistic role of SNPs outside of genic or protein-coding regions on mtDNA: are these variants regulatory? Our analyses indicate statistical associations, which would also be testable by mtDNA editing. These tools would also enable mechanistic investigation of how mtDNA variants impact mitochondrial function (e.g. respiration, proteome), their consequences for cellular processes (e.g. metabolism, epigenome), and how their impact on phenotype and response to diet varies among nuclear backgrounds. We do not dismiss the importance of coding variation, but our data suggest that non-coding variation may yet prove important.

Diet is a major source of biological variation. But the importance of genotype-by-diet variation is increasingly recognized, with genetic variation manifesting phenotypically only under certain dietary conditions, and genotype-specific responses to diet (*1, 24*). We manipulated two specific nutrient classes (EAAs and lipid) normally derived from yeast in fly food, offering greater specificity than previous DMN studies. The nutrients we identify are of particular interest, because EAAs regulate swathes of life-history and health traits, while lipid consumption is associated with the pandemic of human metabolic disease (*3*). We found that impacts of dietary lipid depend on mitonuclear genotype, which may be relevant to understanding variation in impacts of high-fat human diets. The high-EAA diets that we used have parallels to high-protein diets used to increase yields of livestock and human muscle mass, and our finding that EAAs can decrease offspring quality may give pause for thought in use of these diets. We were surprised that EAA enrichment did not enhance offspring development, because we interpreted increased parental egg laying on this food to indicate parental preference, presumably in anticipation of fitness benefits. However, the discrepancy with development and fertility may indicate that EAAs function as signals of food quality as well as metabolites, which could drive deleterious outcomes when EAA levels are not representative of the composition of food that would be found in the yeasts which flies are thought to consume in nature.

An important finding of our study is that mitonucleotype can modify the qualitative output of dietary variation, and can even result in lethality for some genotypes on EAA enriched-diet. For traits where lethality was evident, effect size calculations suggested that impacts of DMN interactions were equal in magnitude to the main effect of diet, suggesting not only that mitonucleotype modulates response to diet, but that DMN interactions can be major sources of variation outright.

We have revealed a relationship between parental nutrition and mitonucleotype. Transient dietary alterations and metabolic disease can drive persistent molecular and phenotypic change, within and across generations(*17, 25*). In our study, transient parental feeding on a high lipid diet even accelerated offspring development in *AA* mitonucleotypes. A direct role for diet in selecting embryos can be excluded because of the standardized post-embryonic environment, and so these effects are likely explained by (A) mitonucleotype-specific parental allocation of development-accelerating factors, after feeding on specific diets, or (B) mitonucleotype-specific selection on offspring from such factors. More generally, after both chronic and parental feeding, effects manifested most strongly in post-embryonic traits, i.e. for fertility, development time, and total progeny, and a dietary and nuclear interactions with C/T polymorphism in *mt:lrRNA* is sufficient to cause these effects. Interestingly, in embryos, lrRNA has been localized outside the mitochondria, in polar granules, suggesting functions in germline determination (*26*), and highlighting this non-coding RNA as a potential mechanistic link to post-embryonic variation. Additional or alternative candidate mechanisms to mediate mitonuclear variation in parental effects include altered epigenetic marks, nutrient provision from mother to offspring, or microbiota. It may be illuminating in the future to ask if effects of parental diet are modulated by parental mitonucleotype (e.g. differential nutrient allocation to eggs, gamete epigenome), or mitonucleotype-dependent processes in offspring (e.g. differential response to altered maternal nutrition). More generally, our study suggests that regulatory variation in mitochondria may modify cellular function in ways that are not yet understood, but appear to depend on dietary and nuclear genetic context. The SNP in lrRNA may, for example, modify protein translation. Altered mitochondrial metabolism will alter overall cellular metabolism, which can have myriad downstream consequences. Much further work is now required to elucidate these mechanisms.

In summary, our study shows that (A) specific nutrients’ fitness effects are determined by interplay of mitochondrial and nuclear genetic variants, (B) DMN effects can produce unprecedented biological variation, and (C) a single-nucleotide substitution in a mtDNA-encoded regulatory factor can underpin these effects. This suggests that regulatory variation in mtDNA may be a hitherto unappreciated but important factor that conspires with nuclear genome to dictate optimal nutrition.

## ACKNOWLEDGMENTS

We thank L Holman, S Parratt, C Selman, E Combet, D Marcu, D Sannino, V Howick and J Rolff for helpful discussion. C Froschauer and R Dobler provided invaluable advice in setting up experiments and the populations.

## FUNDING

UKRI Future Leaders Fellowship MR/S033939/1 (AD)

Dresden Fellowship funded by the Excellence Initiative of the German Federal and State Governments (AD)

University of Glasgow Lord Kelvin Adam Smith Fellowship (AD)

Deutsche Forschungsgemeinschaft Excellence Initiative to TU Dresden (KR)

## AUTHOR CONTRIBUTIONS

Conceptualization: AJD, KR

Methodology: AJD, SV, DKD, KR

Investigation: AJD, SV, LK, LL, EV, RI

Visualization: AJD

Funding acquisition: KR, AJD

Project administration: AJD

Supervision: AJD

Writing – original draft: AJD

Writing – review & editing: AJD, SV, DKD, KR

## DATA AND MATERIALS AVAILABILITY

All data and statistical code will be made available at github.com/dobdobby. Sequence data are being uploaded to NCBI (accession pending).

## SUPPLEMENTARY MATERIALS

**Figure S1.**
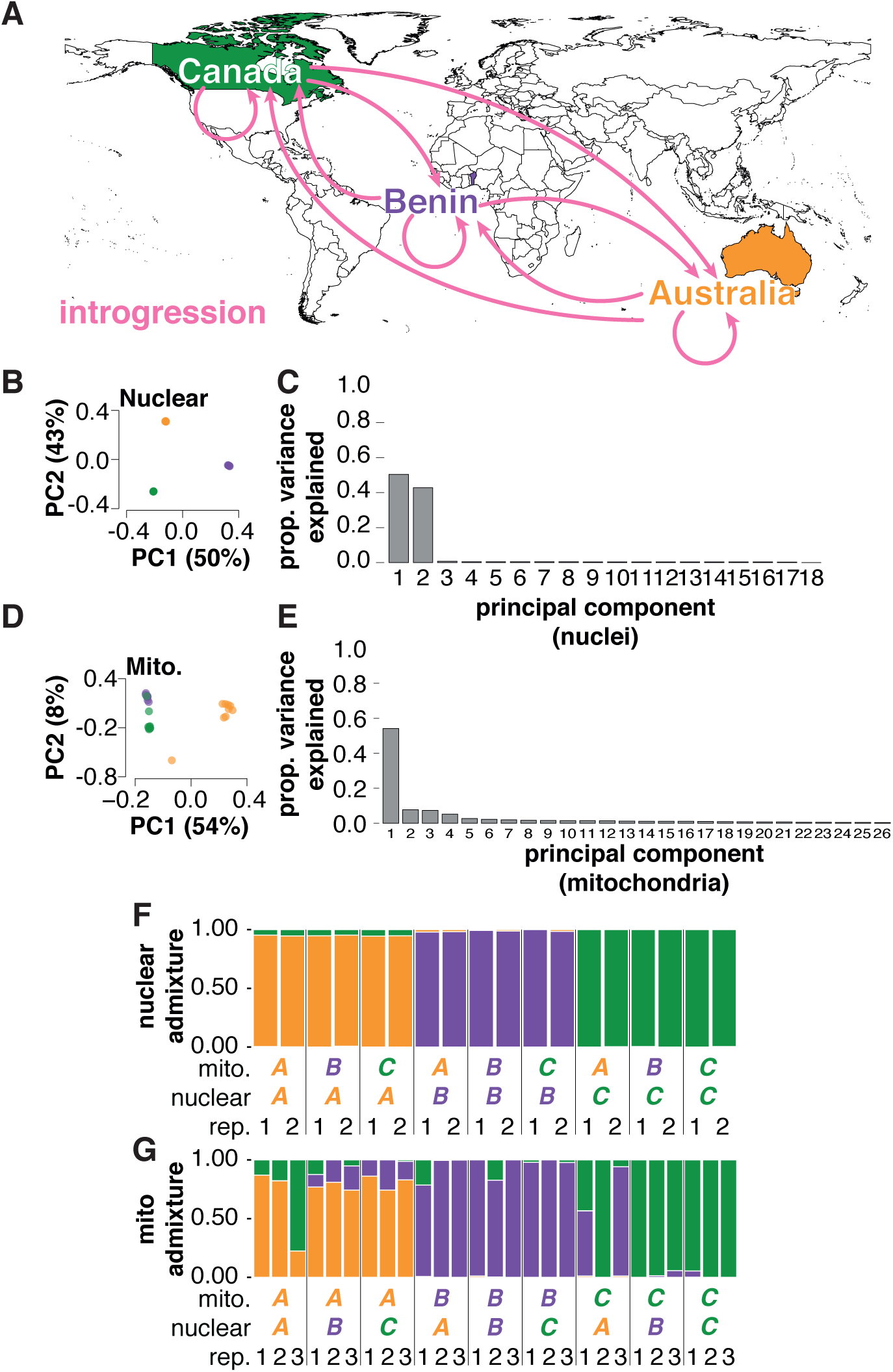
Mitonuclear diversity in a panel of *D. melanogaster*. **A.** Fly populations from Australia, Benin and Canada were introgressed in all possible combinations, generating novel combinations of mitochondrial and nuclear genomes. Three biologically independent replicate populations were established per introgression. **B.** Principal Components Analysis indicates nuclear replacement and homogenization with nuclear genomes from donor populations. PCA was performed on per-population allele frequencies, of all observed nuclear SNPs on the major chromosome arms (2L, 2R, 3L, 3R, and X). Points representing diverse nucleotypes sit on top of one another, suggesting homogenized nuclear genomes even in the presence of distinct mitochondria. **C.** Variance explained by PCA of nuclear SNPs. Barplots show variance explained by each PC. **D.** PCA reveals population grouping by mitochondrial allele frequency, suggesting that the dominant axis of variation (PC1) is driven by SNPs that differentiate mitotypes B and C from mitotype A, with one line (AA_3_) intermediate between the two major clusters. PCA was performed on per-population allele frequencies of observed mitochondrial SNPs. **E.** Variance explained by PCA of mitochondrial SNPs. Barplots show variance explained by each PC. **F.** Nuclear and **G.** mitochondrial admixture proportions. Admixture proportions for each line were inferred by model-based clustering with ConStruct (K=3). To minimize effects of linkage disequilibrium (LD), only nuclear SNPs at least 1kb apart and outside regions of no recombination were considered for PCA and admixture analyses.

**Figure S2.**
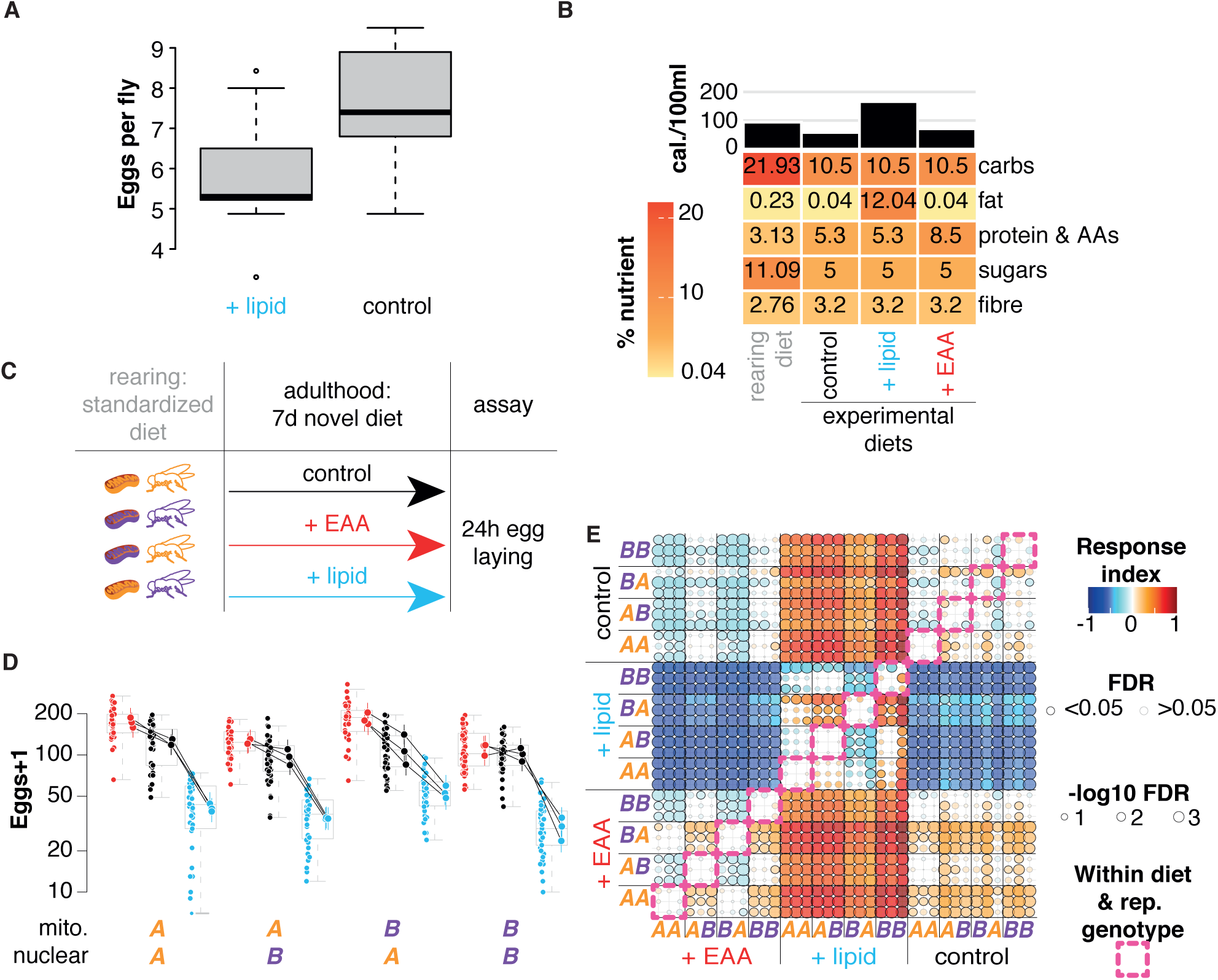
Enriching EAAs or lipid is sufficient to induce repeatable diet-mito-nuclear variation in fecundity. **A.** Reproductive manipulation by enriching fly medium with plant-based lipid. Egg laying by wild-type *Dahomey* flies on control medium (10% yeast, 5% sugar) and control with 15% plant-based lipid source (margarine) added. Boxplots show medians, first and third quartiles and 5th and 95th percentiles. Two-sample t-test t = 1.98, df = 16, p = 0.03. Data shown per fly. **B.** Diet design: The heatmap shows estimated macronutrient content of diets used in this study, bars at top indicate caloric content. **C.** Key & experimental design. Flies were reared from egg to adult on rearing food, and allocated at random to experimental media 6-48h after eclosion, at a density of five of each sex per vial. After seven days, flies laid eggs on fresh food for 24 hours. **D.** Mitonuclear variation in fecundity response to nutrient enrichment. Plot shows eggs laid in vial of five females and five males over 24h. Boxplots show medians, first and third quartiles and 5th and 95th percentiles. Points to left of each box show raw data. Connected points to right of each box show estimated marginal means (EMMs) with 90% confidence intervals. Data shown per vial (5 females + 5 males). **E.** Response to nutritional variation is repeatable among mitonuclear genotypes. Bubble plot shows response index - signed, logged, absolute fold-change in specified comparisons of EMMs - with point size scaled to indicate probability of observed difference (-log10 FDR), and border opacity indicating threshold of statistical significance (FDR≤0.05). Fold-change calculated for conditions on Y-axis relative to conditions on X-axis, e.g. bottom-right cluster of points shows increase on EAA-enriched media relative to control. Points along diagonal show comparisons within replicate genotypes on the same diet, with few significant differences among replicate genotypes. In response to lipid enrichment, the same changes were always evident among replicates of the same genotype, and in response to EAA enrichment similar changes were evident in some replicates. Boxes indicate comparisons among replicates of the same genotype on the same diet.

**Figure S3.**
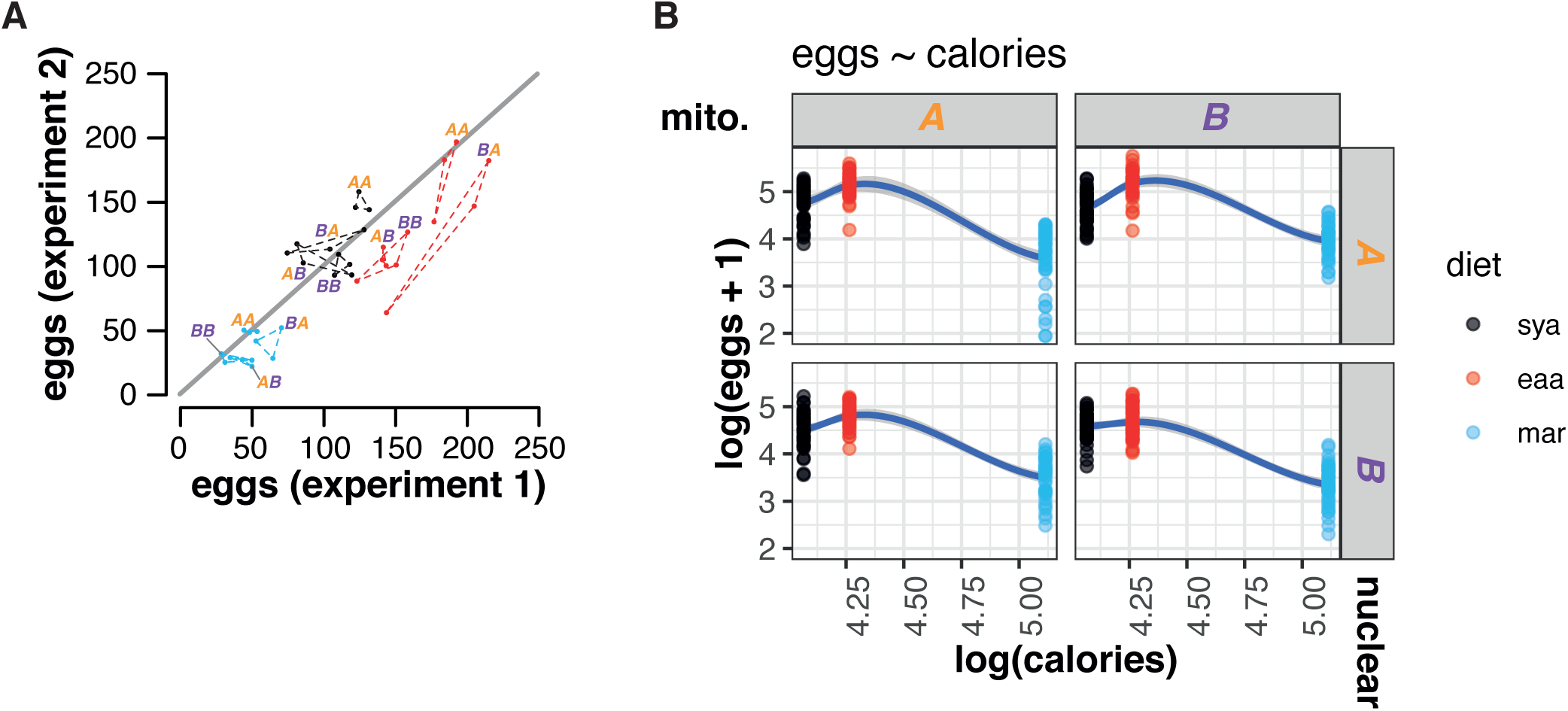
Diet-mito-nuclear variation in fecundity: Technical repeatability and absence of linear relationship to caloric content. **A.** Intra-line correlations in impact of mitonuclear variation on egg laying, and impact of diet on egg laying. Each point shows mean egg laying per line per diet in each of two replicate experiments, with the replicates of each haplotype grouped by dashed lines. Means were correlated between experiments (Pearson’s r = 0.87, p = 7.6e-12). **B.** Fecundity does not correlate caloric content of experimental media. Scatterplots show eggs at each caloric level, with facets per each combination of mitochondrial (columns) and nuclear (rows) genotype. Diet indicated by color. Lines show smoothed spline through points. Trait values do not linearly correlate with calories, therefore caloric content is no more informative than modeling diet as an unordered factor.

**Figure S4.**
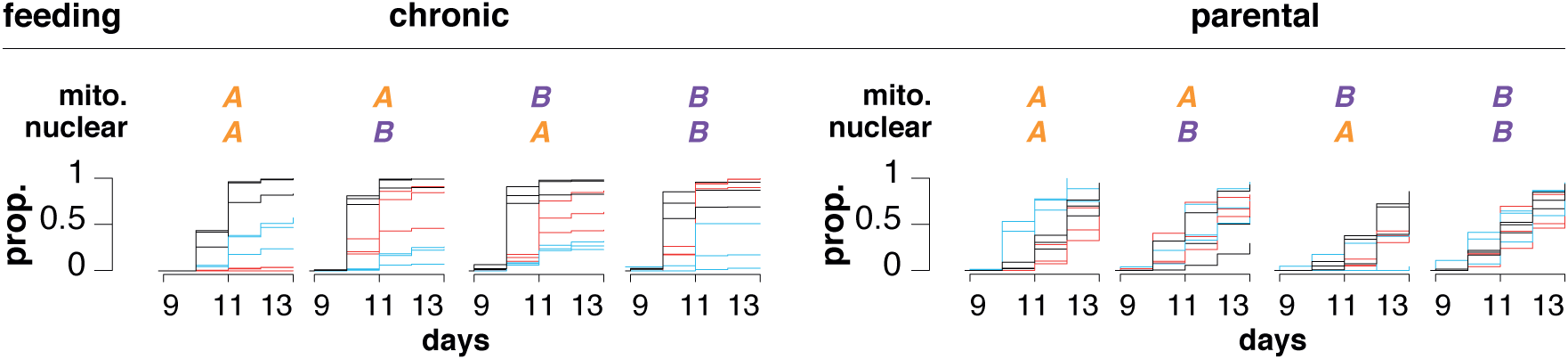
Kaplan-meier plots of development. Plots show proportion eclosed over time. Note the exclusion of EMMs for development of genotype *AA_3_* in chronic feeding: EAA lethality prevented meaningful estimation.

**Figure S5.**
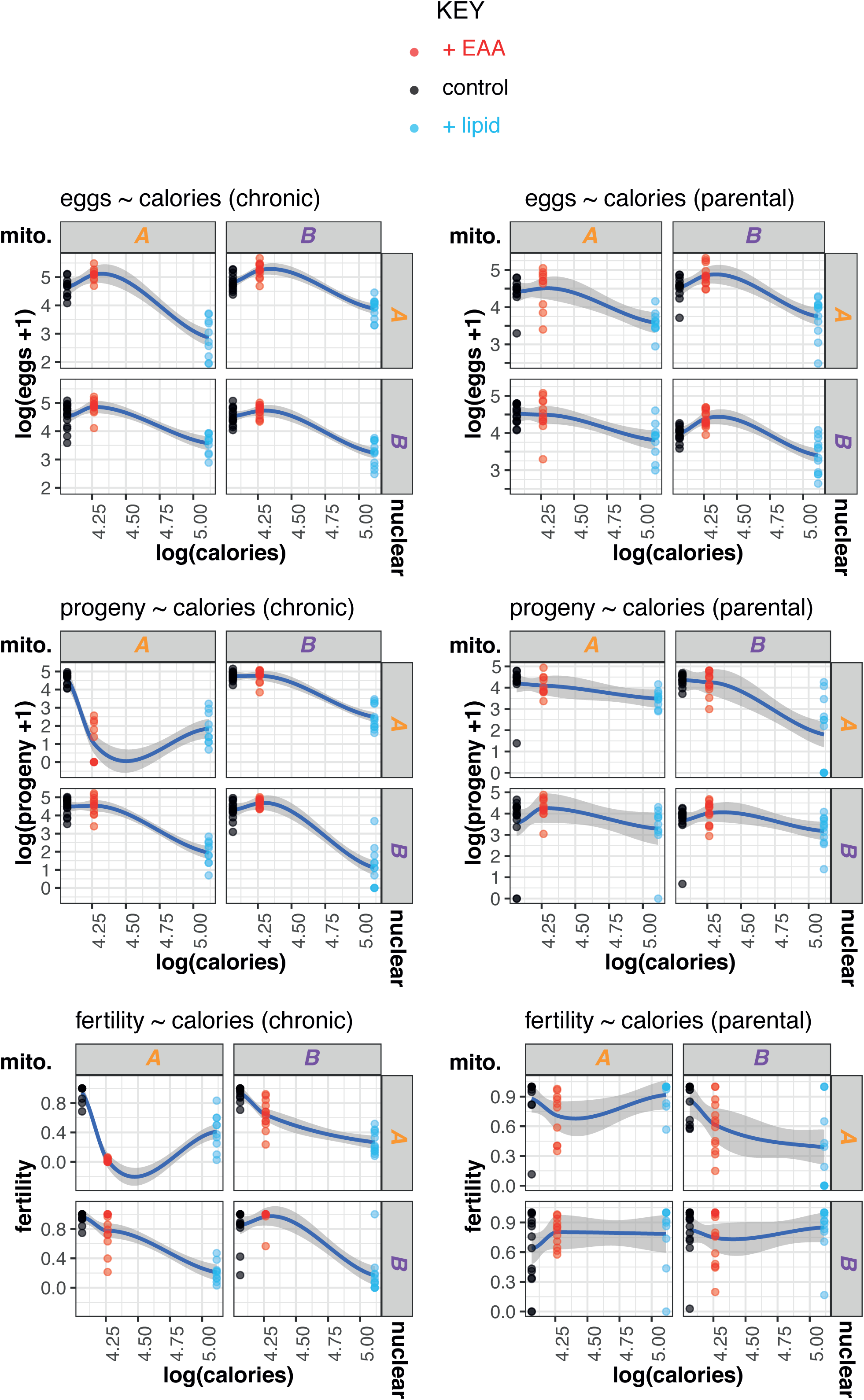
Trait values do not correlate with the caloric content of experimental media. Trait and feeding paradigm indicated above each panel of plots. Within each panel, scatterplots show trait values at each caloric level, with facets per each combination of mitochondrial (columns) and nuclear (rows) genotype. Diet indicated by color. Lines show smoothed spline through points. Trait values do not linearly correlate calories.

**Figure S6.**
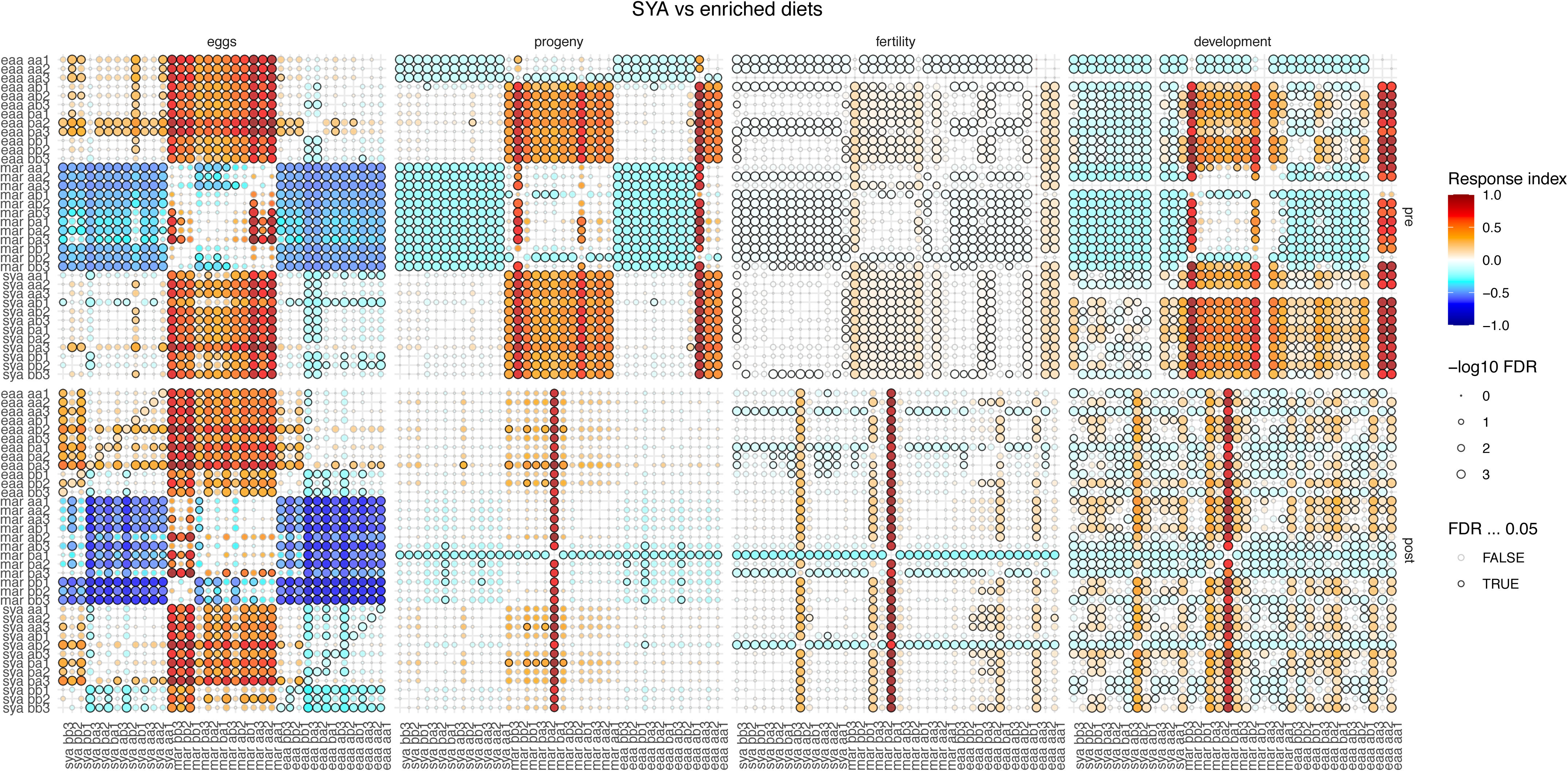
Structure of statistically significant differences among populations. Bubble plot shows response index - signed, logged, absolute fold-change in specified comparisons of EMMs - with point size scaled to indicate probability of observed difference (-log10 FDR), and border opacity indicating threshold of statistical significance (FDR≤0.05). Fold-change calculated for conditions on Y-axis relative to conditions on X-axis, e.g. bottom-right cluster of points shows increase on EAA-enriched media relative to control. Points along diagonal show comparisons within replicate genotypes on the same diet, with few significant differences among replicate genotypes. In response to lipid enrichment, the same changes were always evident in replicate genotypes, and in response to EAA enrichment similar changes were evident in some replicates. Boxes indicate comparisons within replicate populations on the same diet.

**Figure S7.**
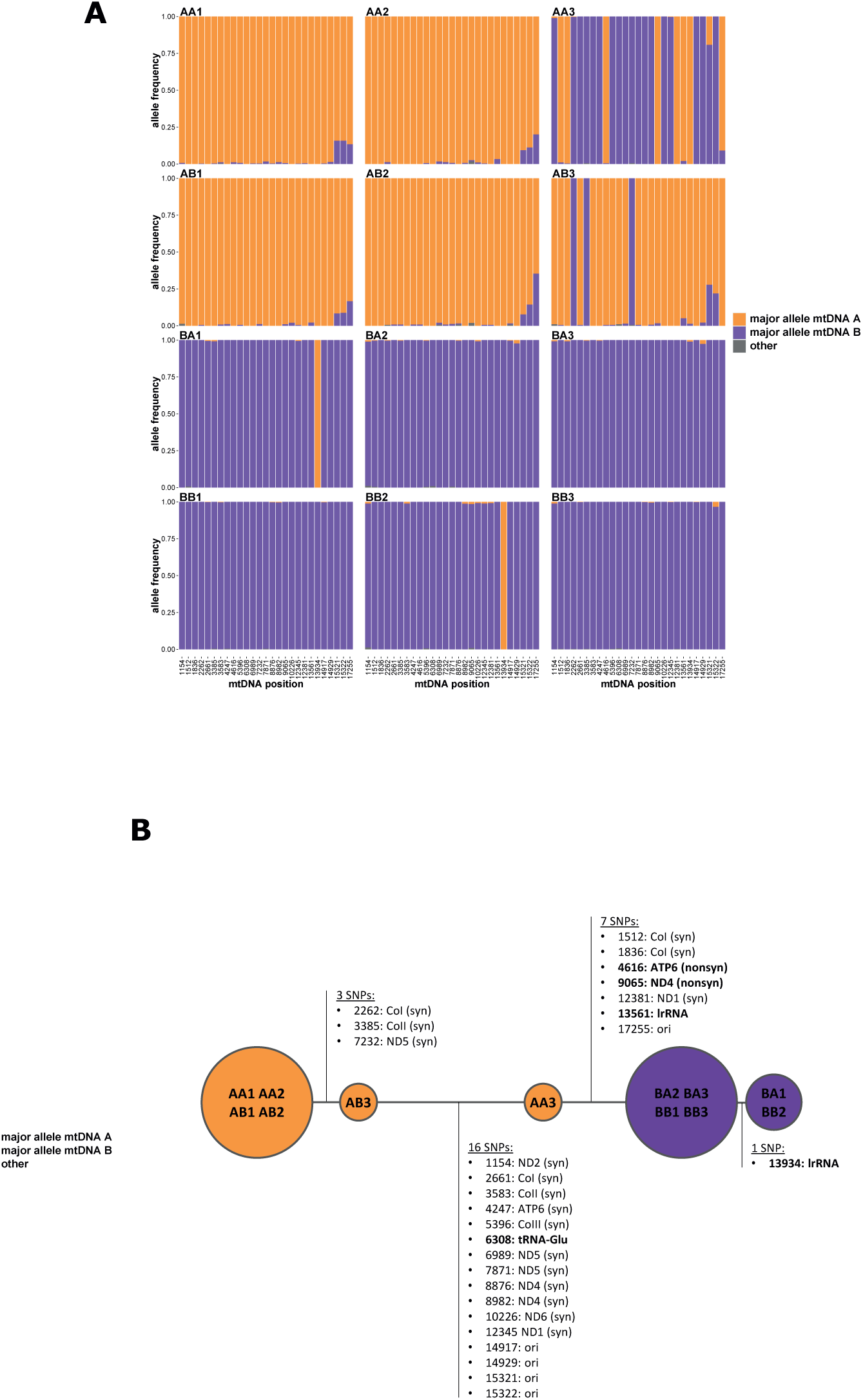
Significantly differentiated loci between mtDNAs *A* and *B*. In total, 146 SNPs were observed within the 12 populations of which 27 were significantly differentiated between populations with mtDNAs of different origins. Significant allele frequency differences were assessed by Fisher’s exact test (FDR<0.001). (A) Allele frequencies and (B) network analysis based on the major alleles in each line at the 27 differentiated sites. Nonsynonymous changes (nonsyn) and nucleotide changes in RNAs are in bold.

**Figure S8.**
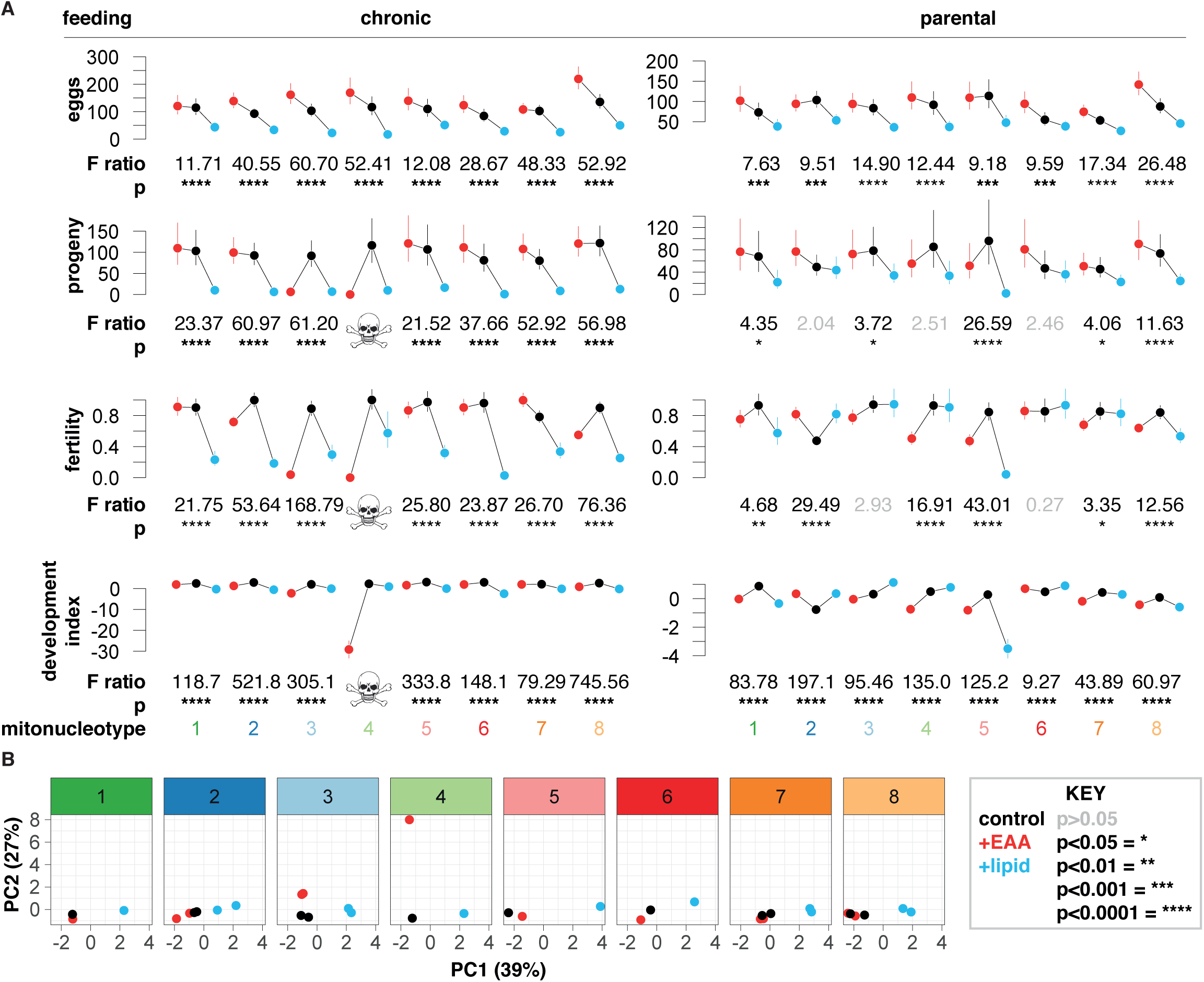
SNP-driven segregation of populations reveals mitonucleotype-specific responses to diet. **A.** Panels show EMMs for traits indicated on y-axes, in feeding conditions indicated at top, with mitonucleotype indicated at bottom. EMMs calculated from GLMMs (fecundity, progeny, fertility) or Cox mixed models (development). Summary statistics beneath each panel were calculated from post-hoc joint tests, showing the impact of diet in each genotype. Skull and crossbone motif indicate exclusion of due to line-specific lethality. Genotype-specific F and P values indicate that non-coding variation in the mitochondrial genome modulates nucleotype-specific response to diet. **B.** PCA shows ordination of populations according to mitonucleotype and diet. PCA values as per Figure 1C, with facets showing mitonucleotypes indicated at top of each panel.

**Table S1.** Rounds of introgression at different phases of the study

| Experiment | N. rounds introgression |
| --- | --- |
| Mitochondrial genome sequencing | 89 |
| Nuclear & mitochondrial genome sequencing | 117 + 120 |
| First fecundity experiment (Fig 2) | 158 |
| Second fecundity experiment (Fig 2) | 160 |
| Fitness indices experiment (Fig 3) | 161 |

**Table S2.** Percentage nuclear admixture (most prevalant background) per population

| **Line** | **% admixture** |
| --- | --- |
| AA1 | 94.9 |
| AA2 | 94.4 |
| AB1 | 97.9 |
| AB2 | 98.3 |
| AC1 | 99.9 |
| AC3 | 99.7 |
| BA2 | 94.8 |
| BA3 | 94.5 |
| BB1 | 99.2 |
| BB3 | 98.7 |
| BC1 | 99.6 |
| BC2 | 99.9 |
| CA2 | 94.5 |
| CA3 | 94.8 |
| CB2 | 99.9 |
| CB3 | 98.5 |
| CC1 | 99.9 |
| CC2 | 99.6 |

**Table S3.**
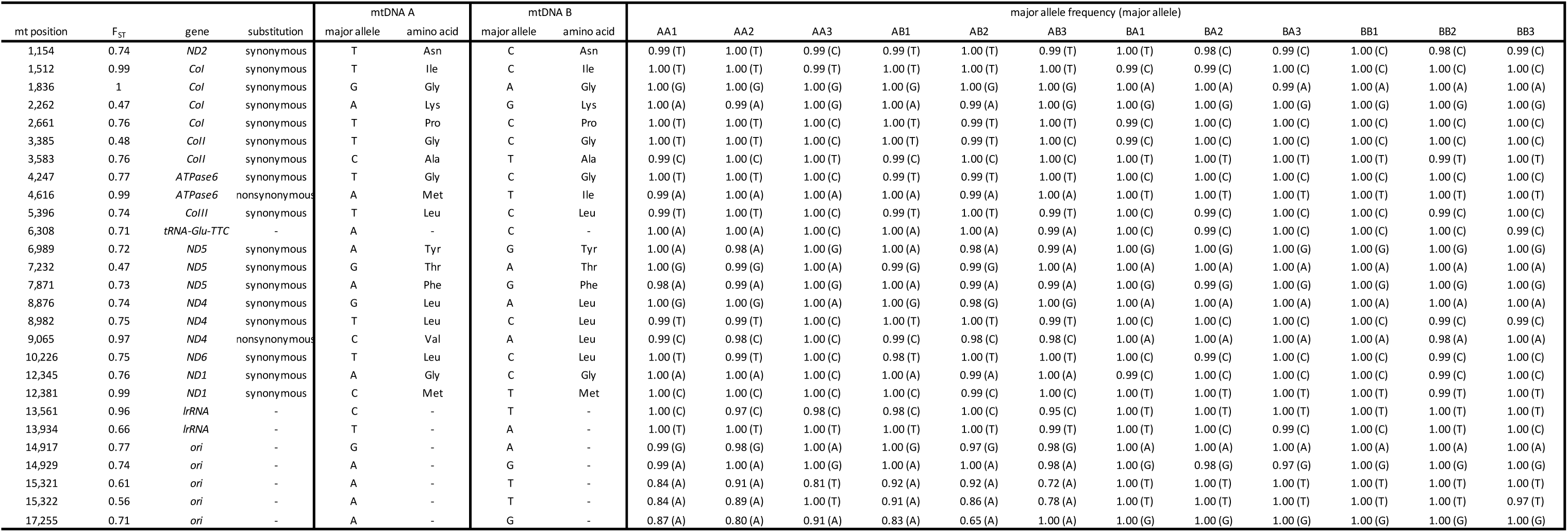
Significantly differentiated SNPs between mtDNAs A and B. SNPs are ranked according to FST. Pairwise FST per SNP were calculated between mitonuclear genotypes (replicates were pooled) based on (64) using Popoolation2 (65). Means of pairwise FST between genotypes with mtDNA A and those with mtDNA B are given. Mitochondrial (mt) positions are according to Flybase Release 6 (66).

| mt position | F<sub>ST</sub> | gene | substitution | mtDNA A |  | mtDNA B |  | major allele frequency (major allele) |  |  |  |  |  |  |  |  |  |  |  |
| --- | --- | --- | --- | --- | --- | --- | --- | --- | --- | --- | --- | --- | --- | --- | --- | --- | --- | --- | --- |
|  |  |  |  | major allele | amino acid | major allele | amino acid | AA1 | AA2 | AA3 | AB1 | AB2 | AB3 | BA1 | BA2 | BA3 | BB1 | BB2 | BB3 |
| 1,154 | 0.74 | ND2 | synonymous | T | Asn | C | Asn | 0.99 (T) | 1.00 (T) | 0.99 (C) | 0.99 (T) | 1.00 (T) | 0.99 (T) | 1.00 (T) | 0.98 (C) | 0.99 (C) | 1.00 (C) | 0.98 (C) | 0.99 (C) |
| 1,512 | 0.99 | Col | synonymous | T | Ile | C | Ile | 1.00 (T) | 1.00 (T) | 0.99 (T) | 1.00 (T) | 1.00 (T) | 1.00 (T) | 0.99 (C) | 0.99 (C) | 1.00 (C) | 1.00 (C) | 1.00 (C) | 1.00 (C) |
| 1,836 | 1 | Col | synonymous | G | Gly | A | Gly | 1.00 (G) | 1.00 (G) | 1.00 (G) | 1.00 (G) | 1.00 (G) | 1.00 (G) | 1.00 (A) | 1.00 (A) | 0.99 (A) | 1.00 (A) | 1.00 (A) | 1.00 (A) |
| 2,262 | 0.47 | Col | synonymous | A | Lys | G | Lys | 1.00 (A) | 0.99 (A) | 1.00 (G) | 1.00 (A) | 0.99 (A) | 1.00 (G) | 1.00 (G) | 1.00 (G) | 1.00 (G) | 1.00 (G) | 1.00 (G) | 1.00 (G) |
| 2,661 | 0.76 | Col | synonymous | T | Pro | C | Pro | 1.00 (T) | 1.00 (T) | 1.00 (C) | 1.00 (T) | 0.99 (T) | 1.00 (T) | 0.99 (C) | 1.00 (C) | 1.00 (C) | 1.00 (C) | 1.00 (C) | 1.00 (C) |
| 3,385 | 0.48 | ColI | synonymous | T | Gly | C | Gly | 1.00 (T) | 1.00 (T) | 1.00 (C) | 1.00 (T) | 0.99 (T) | 1.00 (C) | 0.99 (C) | 1.00 (C) | 1.00 (C) | 1.00 (C) | 1.00 (C) | 1.00 (C) |
| 3,583 | 0.76 | ColI | synonymous | C | Ala | T | Ala | 0.99 (C) | 1.00 (C) | 1.00 (T) | 0.99 (C) | 1.00 (C) | 1.00 (C) | 1.00 (T) | 1.00 (T) | 1.00 (T) | 1.00 (T) | 0.99 (T) | 1.00 (T) |
| 4,247 | 0.77 | ATPase6 | synonymous | T | Gly | C | Gly | 1.00 (T) | 1.00 (T) | 1.00 (C) | 0.99 (T) | 0.99 (T) | 1.00 (T) | 1.00 (C) | 1.00 (C) | 1.00 (C) | 1.00 (C) | 1.00 (C) | 1.00 (C) |
| 4,616 | 0.99 | ATPase6 | nonsynonymous | A | Met | T | Ile | 0.99 (A) | 1.00 (A) | 1.00 (A) | 1.00 (A) | 0.99 (A) | 1.00 (A) | 1.00 (T) | 1.00 (T) | 1.00 (T) | 1.00 (T) | 1.00 (T) | 1.00 (T) |
| 5,396 | 0.74 | ColII | synonymous | T | Leu | C | Leu | 0.99 (T) | 1.00 (T) | 1.00 (C) | 0.99 (T) | 1.00 (T) | 0.99 (T) | 1.00 (C) | 0.99 (C) | 1.00 (C) | 1.00 (C) | 0.99 (C) | 1.00 (C) |
| 6,308 | 0.71 | tRNA-Glu-TTC | - | A | - | C | - | 1.00 (A) | 1.00 (A) | 1.00 (C) | 1.00 (A) | 1.00 (A) | 0.99 (A) | 1.00 (C) | 0.99 (C) | 1.00 (C) | 1.00 (C) | 1.00 (C) | 0.99 (C) |
| 6,989 | 0.72 | ND5 | synonymous | A | Tyr | G | Tyr | 1.00 (A) | 0.98 (A) | 1.00 (G) | 1.00 (A) | 0.98 (A) | 0.99 (A) | 1.00 (G) | 1.00 (G) | 1.00 (G) | 1.00 (G) | 1.00 (G) | 1.00 (G) |
| 7,232 | 0.47 | ND5 | synonymous | G | Thr | A | Thr | 1.00 (G) | 0.99 (G) | 1.00 (A) | 0.99 (G) | 0.99 (G) | 1.00 (A) | 1.00 (A) | 1.00 (A) | 1.00 (A) | 1.00 (A) | 1.00 (A) | 1.00 (A) |
| 7,871 | 0.73 | ND5 | synonymous | A | Phe | G | Phe | 0.98 (A) | 0.99 (A) | 1.00 (G) | 1.00 (A) | 0.99 (A) | 0.99 (A) | 1.00 (G) | 0.99 (G) | 1.00 (G) | 1.00 (G) | 1.00 (G) | 1.00 (G) |
| 8,876 | 0.74 | ND4 | synonymous | G | Leu | A | Leu | 1.00 (G) | 1.00 (G) | 1.00 (A) | 1.00 (G) | 0.98 (G) | 1.00 (G) | 1.00 (A) | 1.00 (A) | 1.00 (A) | 1.00 (A) | 1.00 (A) | 1.00 (A) |
| 8,982 | 0.75 | ND4 | synonymous | T | Leu | C | Leu | 0.99 (T) | 0.99 (T) | 1.00 (C) | 1.00 (T) | 1.00 (T) | 0.99 (T) | 1.00 (C) | 1.00 (C) | 1.00 (C) | 1.00 (C) | 0.99 (C) | 0.99 (C) |
| 9,065 | 0.97 | ND4 | nonsynonymous | C | Val | A | Leu | 0.99 (C) | 0.98 (C) | 1.00 (C) | 0.99 (C) | 0.98 (C) | 0.98 (C) | 1.00 (A) | 1.00 (A) | 1.00 (A) | 1.00 (A) | 0.98 (A) | 1.00 (A) |
| 10,226 | 0.75 | ND6 | synonymous | T | Leu | C | Leu | 1.00 (T) | 0.99 (T) | 1.00 (C) | 0.98 (T) | 1.00 (T) | 1.00 (T) | 1.00 (C) | 0.99 (C) | 1.00 (C) | 1.00 (C) | 0.99 (C) | 1.00 (C) |
| 12,345 | 0.76 | ND1 | synonymous | A | Gly | C | Gly | 1.00 (A) | 1.00 (A) | 1.00 (C) | 1.00 (A) | 0.99 (A) | 1.00 (A) | 0.99 (C) | 1.00 (C) | 1.00 (C) | 1.00 (C) | 0.99 (C) | 1.00 (C) |
| 12,381 | 0.99 | ND1 | synonymous | C | Met | T | Met | 1.00 (C) | 1.00 (C) | 1.00 (C) | 1.00 (C) | 0.99 (C) | 1.00 (C) | 1.00 (T) | 1.00 (T) | 1.00 (T) | 1.00 (T) | 0.99 (T) | 1.00 (T) |
| 13,561 | 0.96 | lrrRNA | - | C | - | T | - | 1.00 (C) | 0.97 (C) | 0.98 (C) | 0.98 (C) | 1.00 (C) | 0.95 (C) | 1.00 (T) | 1.00 (T) | 1.00 (T) | 1.00 (T) | 1.00 (T) | 1.00 (T) |
| 13,934 | 0.66 | lrrRNA | - | T | - | A | - | 1.00 (T) | 1.00 (T) | 1.00 (T) | 1.00 (T) | 1.00 (T) | 0.99 (T) | 1.00 (T) | 1.00 (C) | 0.99 (C) | 1.00 (C) | 1.00 (T) | 1.00 (C) |
| 14,917 | 0.77 | ori | - | G | - | A | - | 0.99 (G) | 0.98 (G) | 1.00 (A) | 1.00 (G) | 0.97 (G) | 0.98 (G) | 1.00 (A) | 1.00 (A) | 1.00 (A) | 1.00 (A) | 1.00 (A) | 1.00 (A) |
| 14,929 | 0.74 | ori | - | A | - | G | - | 0.99 (A) | 1.00 (A) | 1.00 (G) | 1.00 (A) | 1.00 (A) | 0.98 (A) | 1.00 (G) | 0.98 (G) | 0.97 (G) | 1.00 (G) | 1.00 (G) | 1.00 (G) |
| 15,321 | 0.61 | ori | - | A | - | T | - | 0.84 (A) | 0.91 (A) | 0.81 (T) | 0.92 (A) | 0.92 (A) | 0.72 (A) | 1.00 (T) | 1.00 (T) | 1.00 (T) | 1.00 (T) | 1.00 (T) | 1.00 (T) |
| 15,322 | 0.56 | ori | - | A | - | T | - | 0.84 (A) | 0.89 (A) | 1.00 (T) | 0.91 (A) | 0.86 (A) | 0.78 (A) | 1.00 (T) | 1.00 (T) | 1.00 (T) | 1.00 (T) | 1.00 (T) | 0.97 (T) |
| 17,255 | 0.71 | ori | - | A | - | G | - | 0.87 (A) | 0.80 (A) | 0.91 (A) | 0.83 (A) | 0.65 (A) | 1.00 (A) | 1.00 (G) | 1.00 (G) | 1.00 (G) | 1.00 (G) | 1.00 (G) | 1.00 (G) |

**Table S4.** GLMM of fecundity, analysis of deviance table (Type III Wald chi-square tests)

| **Term** | **Chisq** | **Df** | **Pr(>Chisq)** |
| --- | --- | --- | --- |
| (Intercept) | 13063.3133 | 1 | <2.20E-16 |
| mito | 0.0003 | 1 | 0.9854094 |
| nuclear | 69.1324 | 1 | <2.20E-16 |
| diet | 1297.4867 | 2 | <2.20E-16 |
| experi | 33.8005 | 1 | 6.11E-09 |
| mito:nuclear | 3.9543 | 1 | 0.0467519 |
| mito:diet | 1.5993 | 2 | 0.4494799 |
| nuclear:diet | 13.9395 | 2 | 0.0009399 |
| mito:nuclear:diet | 13.8816 | 2 | 0.0009675 |

**Table S5.**
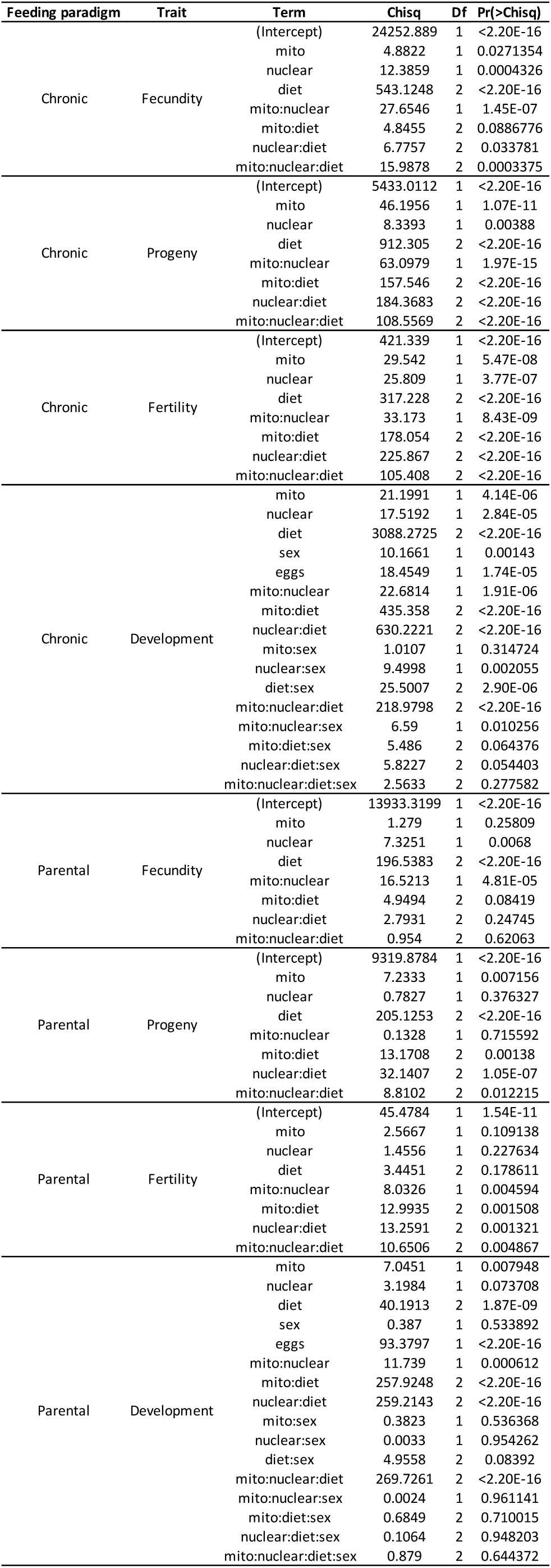
GLMM and CoxME analyses of reproductive traits, analysis of deviance tables (Type III Wald chi-square tests)

**Table S6.**
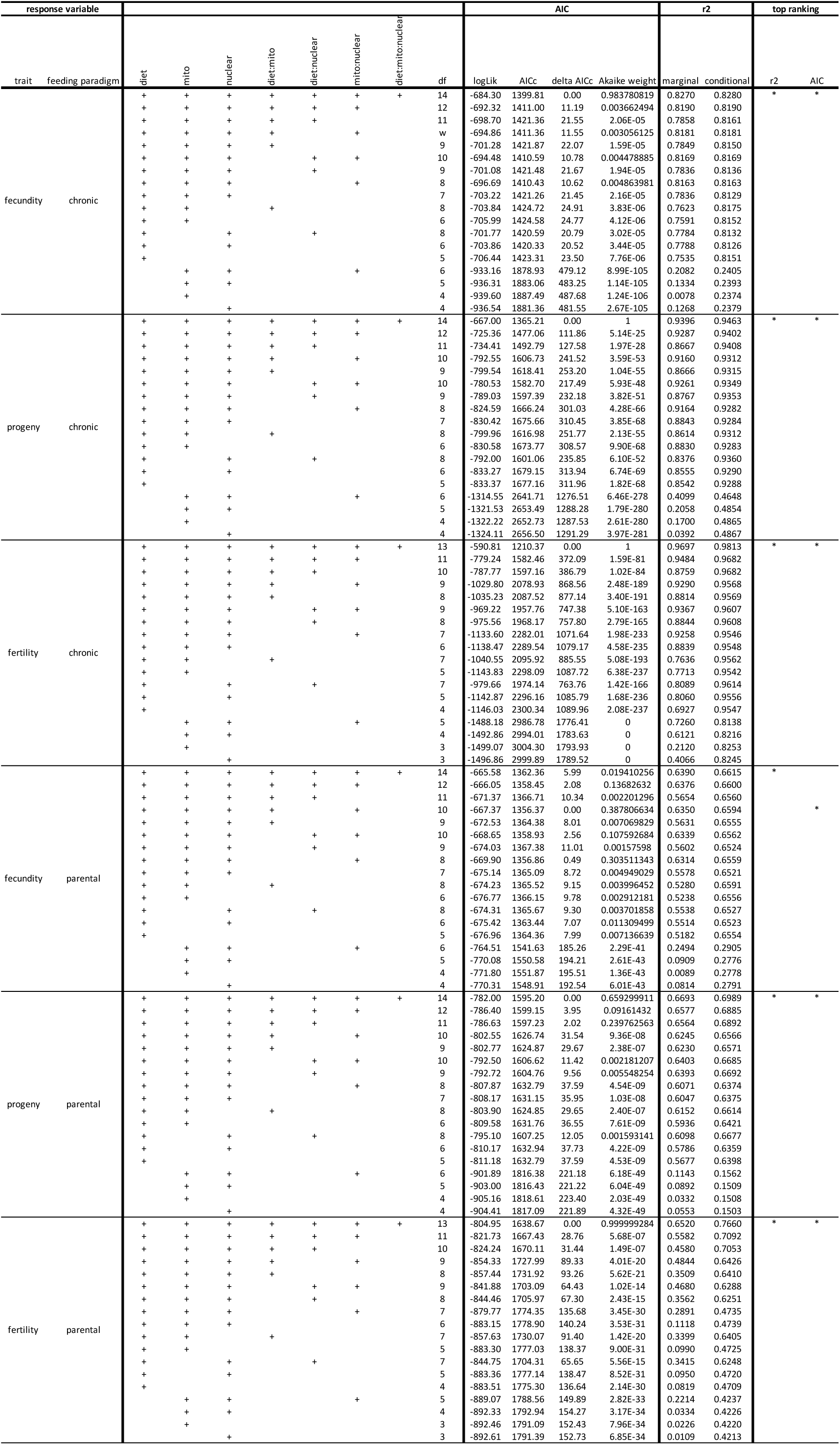
GLMMs of reproductive traits. Model optimisation favors retention of DMN terms. Model terms were systematically included or eliminated, and AICc, delta AICc, Akaike weights and variance explained (r^2) were calculated for each model.

**Table S7.**
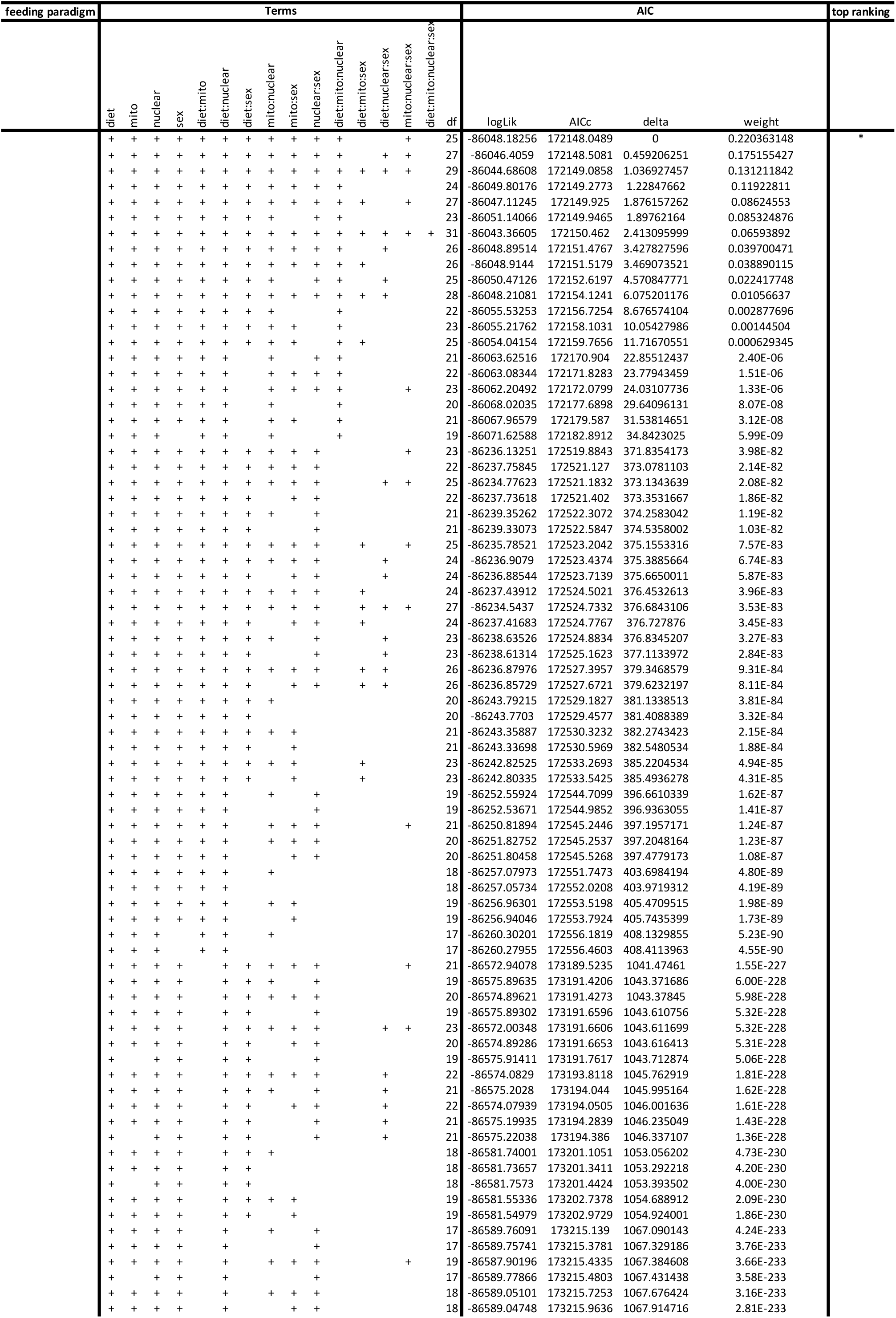
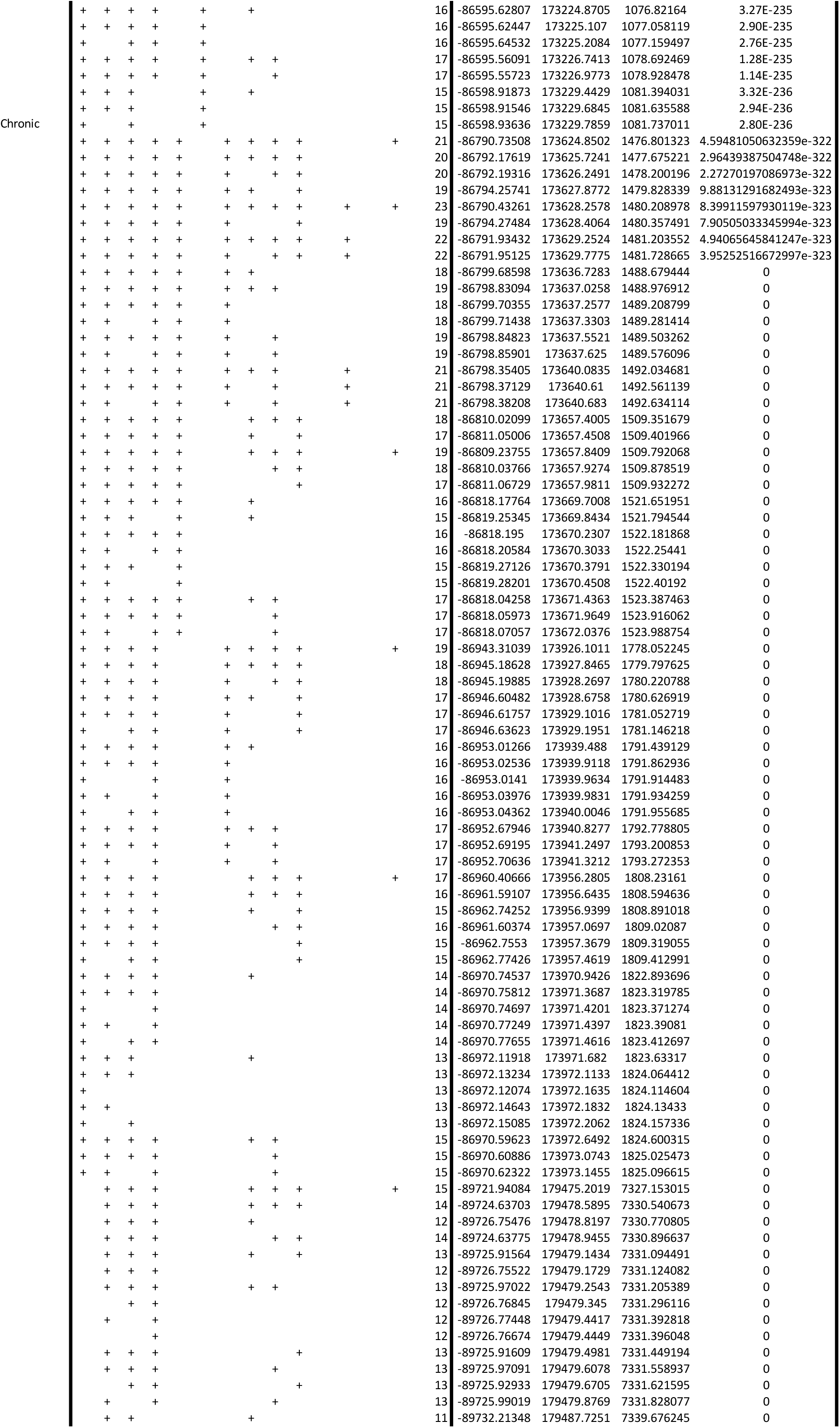
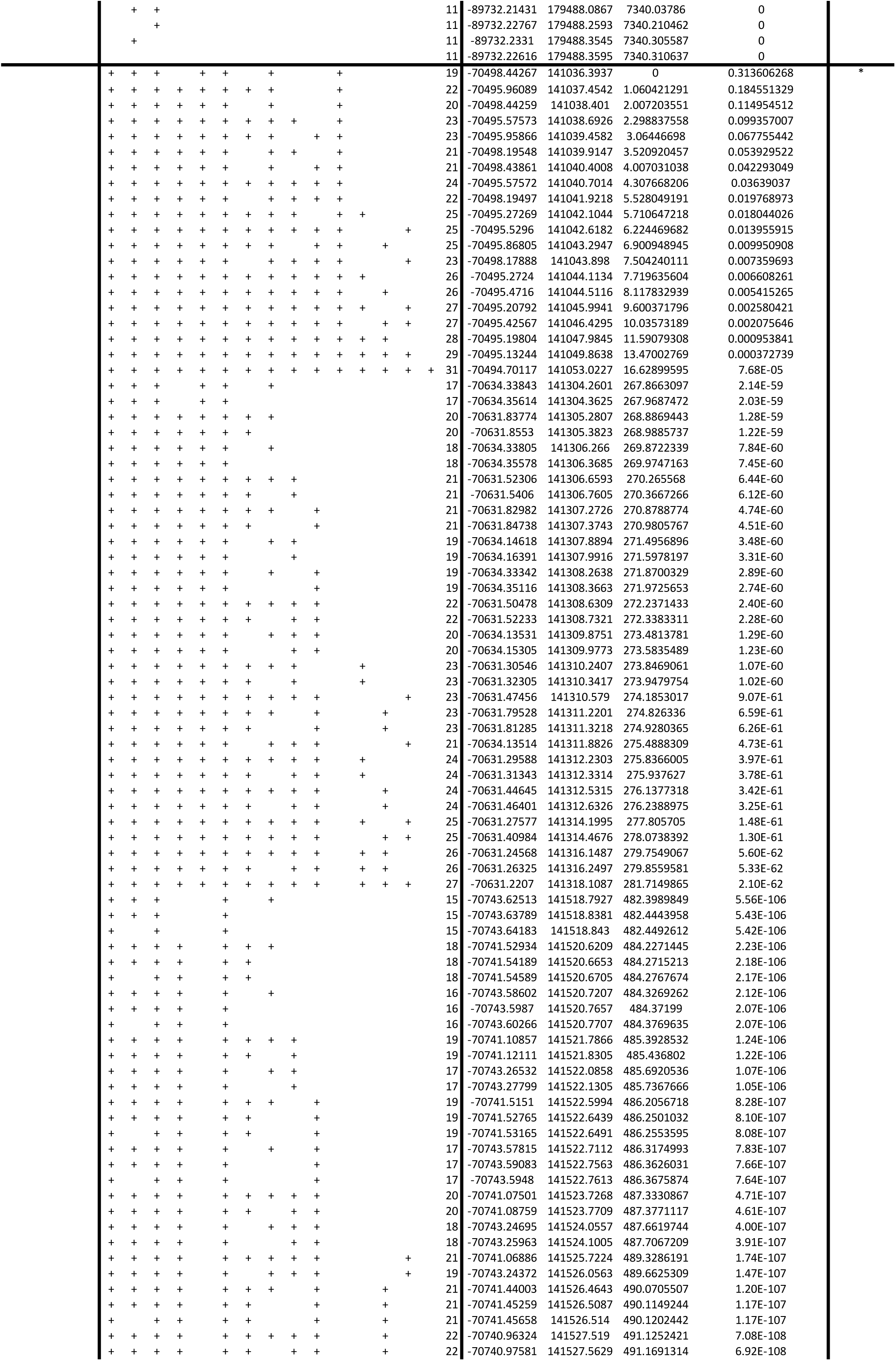
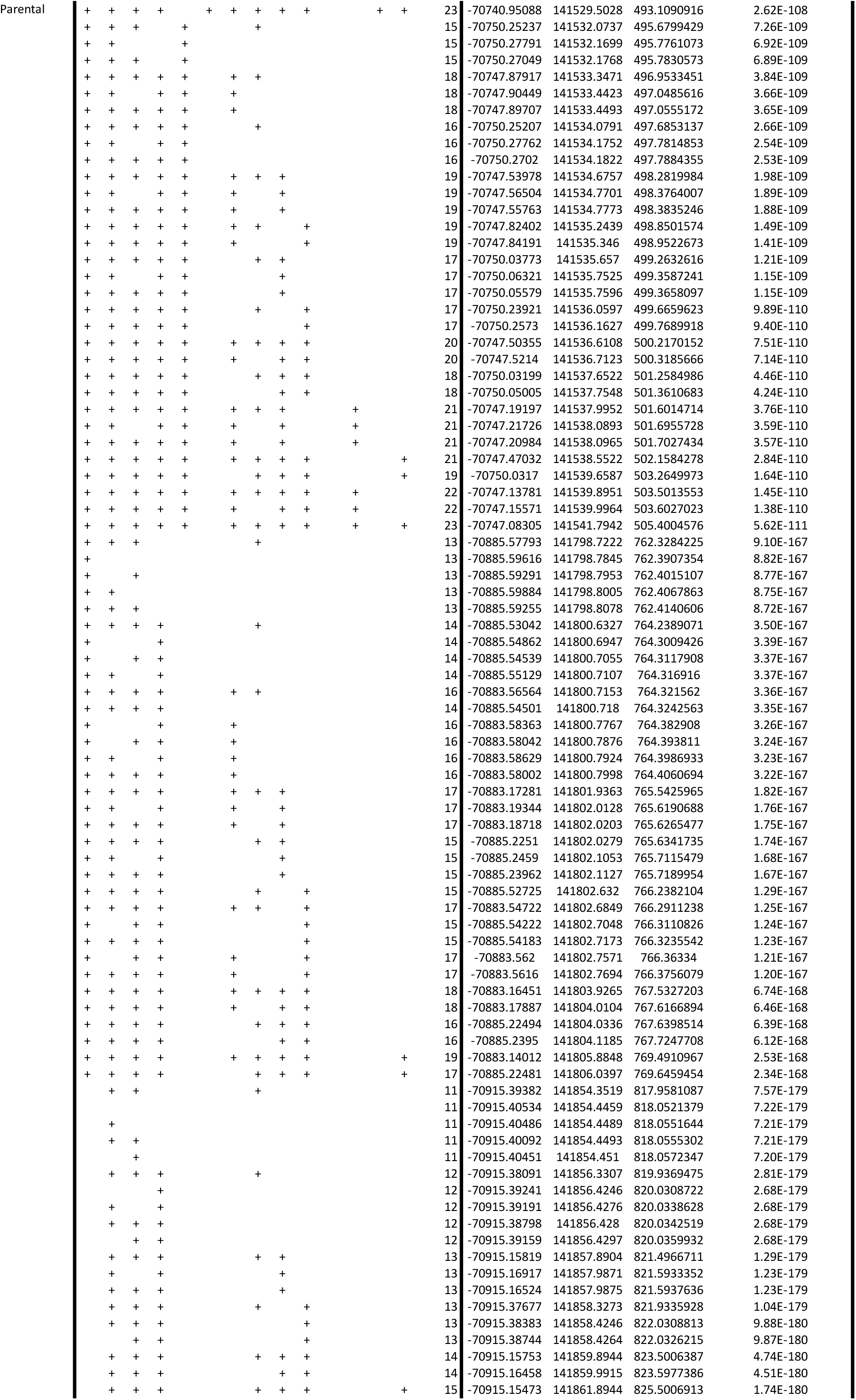
CoxME of development indices. Model optimisation favors retention of DMN terms. Model terms were systematically included or eliminated, and AICc, delta AICc and Akaike weights were calculated for each model.

**Table S8.** mtDNA major allele frequency does not vary substantially over course of introgression. Table shows major allele frequency for indicated mitonucleotypes at indicated times in introgression. Sequence data from 2016 were published by Vaught et al (22). mtDNA from 2018 was collected alongside samples for nuclear DNA sequencing. Allele frequencies for each mitonuclear genotype are based on the two replicates for which sequence data was available in both datasets. Minus (-) correspond to sites with insufficient sequencing depth. Mitochondrial (mt) positions are according to Flybase Release 6.

| mito-nuclear combination |  | AA |  | AB |  | BA |  | BB |
| --- | --- | --- | --- | --- | --- | --- | --- | --- |
| year | 2016 | 2018 | 2016 | 2018 | 2016 | 2018 | 2016 | 2018 |
| mt position |  |  |  |  |  |  |  |  |
| 1154 | 0.995 | 1 | 0.992 | 1 | 0.985 | 0.936 | 0.993 | 0.978 |
| 1154 | 1 | 1 | 1 | 0.993 | 0.996 | 0.941 | 0.999 | 0.992 |
| 1154 | 1 | 1 | 1 | 0.997 | 0.996 | 0.922 | 1 | 0.994 |
| 2262 | 0.994 | 1 | 0.995 | 0.992 | 0.998 | 0.99 | 0.999 | 1 |
| 2661 | 1 | 1 | 0.997 | 0.986 | 1 | 0.913 | 0.998 | 0.985 |
| 3385 | 0.998 | 1 | 0.997 | 0.995 | 0.995 | 1 | 0.997 | 0.986 |
| 3583 | 0.994 | 0.98 | 0.996 | 0.997 | 1 | 0.906 | 0.999 | 0.981 |
| 4247 | 1 | 1 | 0.99 | 0.995 | 0.998 | 0.912 | 0.999 | 1 |
| 4616 | 0.993 | 1 | 0.997 | 1 | 1 | 0.952 | 1 | 1 |
| 5396 | 0.994 | 1 | 0.997 | 0.996 | 0.996 | 0.922 | 0.999 | 0.99 |
| 6308 | 1 | 1 | 1 | 0.996 | 0.996 | 0.898 | 0.997 | 1 |
| 6989 | 0.992 | 1 | 0.991 | 0.995 | 1 | 0.844 | 1 | 1 |
| 7232 | 0.993 | 1 | 0.99 | 0.989 | 1 | 0.984 | 1 | 0.957 |
| 7871 | 0.987 | 1 | 0.995 | 0.992 | 0.997 | 0.872 | 0.998 | 0.971 |
| 8876 | 0.998 | 1 | 0.994 | 0.996 | 0.998 | 0.8 | 0.997 | 1 |
| 8982 | 0.99 | 1 | 1 | 1 | 1 | 0.885 | 0.995 | 1 |
| 9065 | 0.987 | 1 | 0.987 | 1 | 1 | 0.885 | 1 | 1 |
| 10226 | 0.997 | 1 | 0.988 | 0.99 | 0.996 | 0.965 | 1 | 0.987 |
| 12345 | 0.997 | 1 | 0.995 | 0.982 | 1 | 0.886 | 0.999 | 1 |
| 12381 | 0.997 | 1 | 0.998 | 0.997 | 1 | 0.87 | 0.999 | 0.977 |
| 13561 | 0.987 | - | 0.987 | 1 | 1 | 0.881 | 1 | 0.958 |
| 13934 | 1 | 1 | 1 | 0.995 | 0.994 | 0.948 | 1 | 0.989 |
| 14917 | 0.986 | 1 | 0.984 | 1 | 1 | 0.913 | 1 | 1 |
| 14929 | 0.991 | 1 | 1 | 1 | 0.976 | 0.914 | 1 | 1 |
| 15321 | 0.887 | - | 0.919 | 1 | 1 | 0.963 | 1 | 1 |
| 15322 | 0.875 | - | 0.892 | 0.964 | 1 | 0.938 | 0.979 | 1 |
| 17255 | 0.844 | - | 0.724 | - | 1 | - | 1 | - |

**Table S9.** GLM and CoxPH analyses of reproductive traits, including sequence-driven mitonucleotype, analysis of deviance tables (Type III Wald chi-square tests)

| Feeding paradigm | Trait | Term | Chisq | Df | Pr(>Chisq) |
| --- | --- | --- | --- | --- | --- |
| Chronic | Fecundity | diet | 464.94 | 2 | <2.20E-16 |
|  |  | mitoNuclear | 64.53 | 7 | 1.87E-11 |
|  |  | diet:mitoNuclear | 38.52 | 14 | 0.0004326 |
| Chronic | Progeny | diet | 465.93 | 2 | <2.20E-16 |
|  |  | mitoNuclear | 157.37 | 7 | <2.20E-16 |
|  |  | diet:mitoNuclear | 230.07 | 14 | <2.20E-16 |
| Chronic | Fertility | diet | 1005.33 | 2 | <2.20E-16 |
|  |  | mitoNuclear | 291.03 | 7 | <2.20E-16 |
|  |  | diet:mitoNuclear | 1205.51 | 14 | <2.20E-16 |
| Chronic | Development | mitoNuclear | 695.4 | 7 | <2.20E-16 |
|  |  | diet | 5480.1 | 2 | <2.20E-16 |
|  |  | sex | 0 | 1 | 1 |
|  |  | eggs | 23.1 | 1 | 1.52E-06 |
|  |  | mitoNuclear:diet | 2035.6 | 14 | <2.20E-16 |
|  |  | mitoNuclear:sex | 9.2 | 7 | 0.24108 |
|  |  | diet:sex | 6.3 | 2 | 0.04274 |
|  |  | mitoNuclear:diet:sex | 15.5 | 14 | 0.34802 |
| Parental | Fecundity | diet | 166.224 | 2 | <2.20E-16 |
|  |  | mitoNuclear | 51.815 | 7 | 6.35E-09 |
|  |  | diet:mitoNuclear | 16.571 | 14 | 0.2797 |
| Parental | Progeny | diet | 77.94 | 2 | <2.20E-16 |
|  |  | mitoNuclear | 23.689 | 7 | 0.001292 |
|  |  | diet:mitoNuclear | 45.666 | 14 | 3.18E-05 |
| Parental | Fertility | diet | 67.008 | 2 | 2.81E-15 |
|  |  | mitoNuclear | 187.741 | 7 | <2.20E-16 |
|  |  | diet:mitoNuclear | 278.077 | 14 | <2.20E-16 |
| Parental | Development | mitoNuclear | 762.38 | 7 | <2.20E-16 |
|  |  | diet | 240.77 | 2 | <2.20E-16 |
|  |  | sex | 2.91 | 1 | 0.087805 |
|  |  | eggsD2 | 149.48 | 1 | <2.20E-16 |
|  |  | mitoNuclear:diet | 1523.17 | 14 | <2.20E-16 |
|  |  | mitoNuclear:sex | 7.42 | 7 | 0.386128 |
|  |  | diet:sex | 9.66 | 2 | 0.007998 |
|  |  | mitoNuclear:diet:sex | 19.14 | 14 | 0.159726 |

**Table S10.**
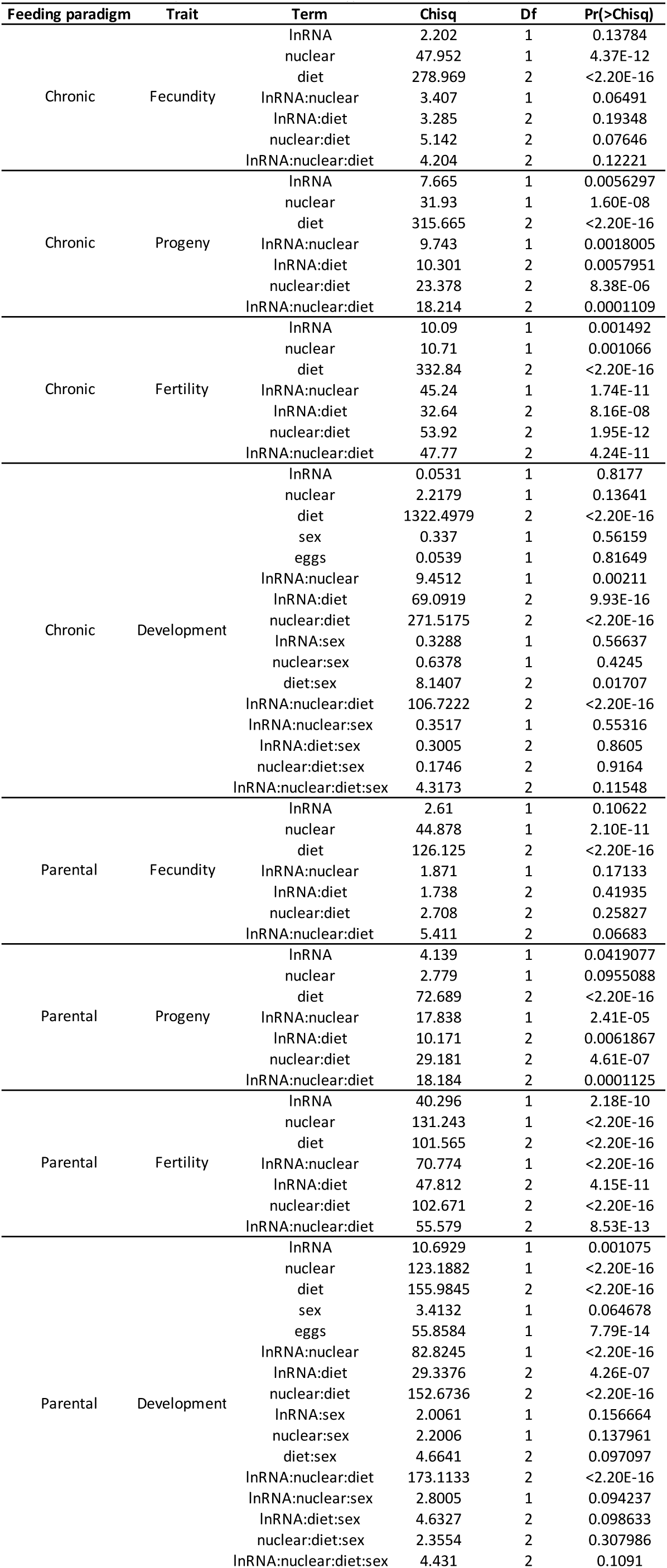
GLM and CoxPH analyses of reproductive traits, in flies where only known mtDNA SNP is in mt:lrRNA. Analysis of deviance tables (Type III Wald chi-square tests)

### Supplementary text

We produced *D. melanogaster* populations (Figure S1A) comprising replicated and fully-factorial combinations of mitochondrial and nuclear genomes from Australia, Benin and Canada (*A*, *B*, *C*, respectively). The crossing scheme was designed to produce distinct mitochondrial backgrounds bearing equivalent pools of standing nuclear variation, by introgressing populations either reciprocally or to themselves. For brevity, we abbreviate population names, giving mitochondrial and then nuclear origin (e.g. *AB* = Australian mitochondria, Beninese nuclei). Each combination was triplicated (e.g. *AB_1_, AB_2_*, *AB_3_*) at the beginning of the introgression and maintained in parallel throughout more than 160 introgressions, altogether generating 27 populations, comprising nine triplicated mitonucleotypes.

After >100 introgressions (Table S1), assuming no mitochondrial incompatibility, we expected nucleotypes to be replaced, and that variation among population with co-originating nuclear backgrounds would be indistinguishable, regardless of mitotype. For a proof of principle, single nucleotide polymorphisms (SNPs) in the nuclear genome were identified by Pool-seq, sampling two populations per mitonucleotype (i.e. 18/27 total). Within each nucleotype, principal components analysis (PCA) suggested negligible differentiation by mitotype, with all samples in each nucleotype sitting indistinguishably on top of one another in an ordination plot (Figure S1B), on axes that explained 93% total variance (Figure S1F). Admixture analysis found between 94.4% and 99.9% replacement of nuclear genomes (Figure S1F, Table S2). These analyses confirmed that the panel of populations was suitable for studying the behaviors of distinct mitotypes in populations with comparable nuclear genomic diversity.

We reanalyzed previously-reported mitochondrial sequences for the populations (*1*), finding 28 significantly differentiated SNPs (Fisher’s exact test, FDR<0.001). PCA (Figure S1D) revealed consistent within-mitotype clustering, except population *AA_3_*, which was distinct from other *A* mitotypes. Little differentiation was apparent between mitotypes *B* and *C* on the PCA, and only one SNP was significantly differentiated between these mitotypes. These findings suggested that phenotyping both *B* and *C* mitochondria in the same experiment would likely prove redundant. By contrast, 27 of the 28 SNPs were significantly differentiated between mitotypes *A* and *B* (Table S3), i.e. the majority of mitochondrial diversity in the full panel was represented by these two mitotypes. Therefore, we chose to work with only one mitochondrial background, retaining Beninese mitochondria because of the widespread use of this background in fly research, and excluding Canadian mitochondria (i.e. studying only *AA, AB, BA* and *BB*). We also excluded Canadian nucleotypes (*AC, BC*) because of lack of coevolutionary history with *A* or *B* mitotypes. This reduced number of populations for phenotyping from 27 to 12.

#### Novel high-lipid diet represses fecundity

Previous studies showed mitonuclear modulation of effects of diluting dietary yeast (*2–6*), which is a source of multiple nutrients. We studied effects of specific nutrients, normally derived from yeast, by specifically enriching either lipids or essential amino acids (EAAs). EAA enrichment increases fly fecundity ∼20% (*7–9*). We were interested in lipids because of relevance to Western human diets (*10, 11*), and so we developed a new high-lipid diet (see materials & methods). Lipid decreased wild-type fecundity by 20% (Figure S2A), confirming that this diet modulates reproduction, but inversely mirroring EAA’s effect. Figure S2B shows approximate nutrient contents of control, EAA-enriched and lipid-enriched diets. Flies were maintained on a distinct “development” medium prior to experiments, to distribute any novelty effects of switch to experimental food evenly among conditions.

#### Specific nutrients sufficient for DMN variation

After ≥158 introgressions of the flies, EAA and lipid were each sufficient to cause DMN variation in reproduction (Figure S2B, Figure S2C). Repeating the same assay across multiple generations of introgression gave the same results, without differences that would indicate intergenerational drift (Figure S3A). Fecundity did not correlate with the increased caloric density in the EAA and lipid enriched diets (Figure S3B), as shown previously (*12, 13*): fecundity therefore cannot be explained as a simple linear effect of calories. Diet was therefore modelled as an unordered factor. A DMN interaction was visually apparent (Figure S2D). A statistical model (GLMM, Table S4), which explained the majority of variance (conditional R^2^=0.768), confirmed the DMN interaction (p=0.001).

To assess repeatability among genetic replicates, and to identify differential effects of EAA versus lipid enrichment, we calculated estimated marginal means (EMMs) (*14*) for each line on each diet. EMMs were visually correlated among genetic replicates (Figure S2D). To assess repeatability, we calculated an index of differences between conditions (Supplementary text) and p-value for each pairwise diet:line comparison (Figure S2E). First, we used these values to assess consistency among the genetic replicates of each mitonucleotype (e.g. *AA_1_*, *AA_2_*, *AA_1_*) which would be refuted by frequent among-replicate differences on the same diet. Of the 36 total comparisons (3 diets, 12 populations), only four were significantly different, indicating that mitonuclear replication led to repeatable fecundity. Second, we assessed whether genetic replicates responded equivalently to dietary change. Adding lipid consistently repressed fecundity, but the magnitude varied repeatably by mitonucleotype (Figure S2D, FIgure S2E). Responses to EAA enrichment were more nuanced but were still overall repeatable among replicates, with most populations increasing egg laying, to the greatest extent in *BA* populations but more modestly in *AA*; and fewer populations responding to EAAs in *B* nucleotypes independent of mitochondria (Figure S2D, Figure S2E). These analyses confirmed that (A) fecundity across varied diets was repeatable among independently replicated mitonucleotypes, and (B) dietary EAAs and lipid can cause this variation

#### AIC and r^2^ calculations

For orthogonal tests of the importance of DMN terms, we asked if alternative models, excluding DMN terms, were better descriptors of the data. We fit a structured series of models for each trait, systematically including or eliminating diet, mitotype and nucleotype, and their interactions. Interactions with offspring sex were also included for development models. We calculated Akaike Weights (Table S6, Table S7), which evaluate the relative performance of a set of models. For all traits but one (fecundity, in parental feeding paradigm), Akaike Weights showed that the best-performing models included DMN interaction terms. For development time in each feeding paradigm, higher interactions between sex and diet, mitotype and nucleotype were generally not favored, consistent with the suggestion from effect size calculations that offspring sex was not a major modulator of genetic and dietary effects in these flies.

We also calculated variance explained (r^2^) by GLMMs (fecundity, progeny, fertility: r^2^ cannot be calculated for Cox models of development time), to evaluate each model’s capacity to statistically predict phenotype. GLMMs favored by Akaike Weights explained between 65% and 95% of variance (Table S6), suggesting that most sources of variation in our flies were accounted for by our models.

#### Mitochondrial SNP analysis

All 27 SNPs in *A* and *B* populations were biallelic (Figure S7A). 70% of major alleles were fixed, and 89% were at frequency ≥0.99 (Figure S7A, Table S3). We mapped the mitochondrial SNPs to genes and regulatory sequences. 19/27 mitochondrial SNPs were in protein-coding regions, in genes encoding subunits of electron transport chain complexes and *ATPase subunit 6,* but only 2/19 were predicted to be nonsynonymous (in *ATPase subunit 6* and *ND4*). However, 8/27 SNPs were in non-coding regions (in the origin of replication, a tRNA, and *long ribosomal RNA*). Thus, the majority of significantly differentiated SNPs in our populations do not appear to change amino acid sequence. Resequencing at a two-year interval indicated that these frequencies were stable over time and unlikely to have changed substantially between the time of genotyping and phenotyping (Table S8), suggesting it was valid to associate this mitochondrial genetic variation with phenotype.

We examined among-line distribution of major mitochondrial alleles. Network analysis (Figure S7B) revealed a punctuated continuum of among-line variation, independent of nucleotype, including some populations with unique mitotypes (*AB_3_, AA_3_*), but others with identical mitotypes, even over distinct nucleotypes (*AA_1_, AA_2_, AB_2_, AB_2_*; and *BA_2_, BA_3_, BB_1_, BB_3_*; and *BA_1_, BB_2_*). This analysis, focussed just on populations phenotyped in this study, reinforced findings of our previous network analysis of the full set of populations (shown in Figure S1) (*1*).

#### Associations between mito-nucleotype and differential responses to diet

We studied the interaction between diet and sequence-driven mitonucleotype. We used mitonucleotype rather than SNPs because (A) tests of linked SNPs would have been redundant; and (B) *AA_3_*’s mitotype did not occur over the *B* nucleotype, so its interaction with nucleotype could not be assessed.

Re-calculating and plotting EMMs per mitonucleotype indicated considerable variation in response to diet (Figure 2B). We excluded mitonucleotype 4 (line *AA_3_*) from statistical analysis because its extreme trait values complicated modelling. We do not discard this line as an outlier: rather we highlight its extreme and unique response to diet, which is self-evident without statistical modelling. Among other populations, ANOVA tests revealed significant mitonucleotype:diet interactions (Table S9). To estimate variability in response to dietary change, we calculated F-ratios and P-values for effect of diet per mitonucleotype. F-ratios varied up to 10-fold, depending on trait. Diet effects were significant for all mitonucleotypes in the chronic feeding paradigm (p<0.001 in all cases), but not in the parental feeding paradigm. These analyses suggest that variance in response to diet can be partitioned by sequence-based mitonucleotype.

### Supplementary text - Materials & methods

#### Diets

Development medium contained 1.4% agar and 4.5% brewer’s yeast (both Gewürzmühle Brecht, Germany), 10% cornmeal and 11.1% sucrose (both Mühle Milzitz, Germany) (all w/v), 0.45% propionic acid and 3% nipagin (v/v). Experimental media built on published protocols (*8, 9, 15*). These media contained a final concentration of 10% brewer’s yeast, 5% sucrose, 1.5% agar (w/v), 3% nipagin and 0.3% propionic acid (v/v). EAAs were purchased as powder (Sigma), and supplemented by dissolving into a 6.66x solution in ddH_2_0 pH 4.5 (final media concentrations: L-arginine 0.43 g/l, L-histidine 0.21 g/l, L-isoleucine 0.34 g/l, L-leucine 0.48 g/l, L-lysine 0.52 g/l, L-methionine 0.1 g/l, L-phenylalanine 0.26 g/l, L-threonine 0.37 g/l, L-tryptophan 0.09 g/l, L-valine 0.4 g/l). We added margarine (15% w/v, after (*16*)) to ensure that lipid supply was plant-based, because wild fly physiology is likely influenced by their consumption of what appears to be a vegan diet (*17, 18*), and because margarine sets in agar (in contrast to oils). Margarine (*Ja! Pflanzenmargarine* from Rewe Supermarkets, Germany; manufacturer’s analysis 720 kcal / 100g; 80/100g fat from 23/100g saturated fatty acids, 40/100g monounsaturated fatty acids, 17/100g polyunsaturated fatty acids) was briefly melted then mixed thoroughly into the food (15% w/v). Final nutrient contents of rearing and control media were estimated using the *Drosophila* diet content calculator (*19*), with additional protein, lipid and caloric content after nutrient supplements calculated according to margarine nutrient content report, and by assuming caloric equity between EAAs and protein at a caloric value of 4 calories/g (USDA). Vials contained ∼5ml of food, and were stored at 4°C for up to 1 week before use.

#### Flies

*D. melanogaster* fly lines were established as described in Figure S1 and maintained on development medium throughout their history prior to our experiments. The ancestral Australian population was isolated in Coffs Harbour, NSW, Australia (*20*). The Benin population is the widely-used *Dahomey* population, isolated in the 1970s in Dahomey (now Benin). The cytoplasmic endosymbiont *Wolbachia* was cleared by tetracycline treatment 66 generations prior to experiments. For each line, 45 females of the desired mitochondrial background were crossed to 45 males of the desired nuclear background per generation. Iterating this process over many generations led to introgression of the desired nuclear background (from males) into each mitochondrial background. Fly lines were maintained at 25°C on development medium throughout their history prior to experimentation. For experiments, flies were collected upon eclosion to adulthood and fed fresh developmental medium, before being pooled, split and assigned at random to experimental medium in groups of 5 males and 5 females. Experimental flies were maintained at 25°C, and transferred to fresh media every 48-72h for one week. Flies were transferred to fresh medium 24h before egg laying experiments. For development experiments, eggs were incubated at 25°C and pupation and eclosion were scored daily. Eclosing adults were lightly CO_2_- anaesthetised before counting and sexing.

#### Genome sequencing

Input DNA controls from an unpublished ChIP-Seq experiment were used for whole genome sequence analysis. Genomes of two replicates for each mitonuclear combination were sequenced. Pools of 50 adult flies were subjected to a standard native ChIP protocol as described in Cosseau *et al.* 2009. The protocol included an MNase digestion step of six minutes at 37°C using 15U of the enzyme (Thermo Fisher Scientific) per sample which yielded fragments between 284 and 300bp in length. DNA was extracted with the QIAquick PCR purification kit (Qiagen) and 100ng genomic DNA of the unChIPped input/negative controls were used for library preparations with NEB Next Ultra DNA lib Prep kit for Illumina. Over 50 million 2×75bp (PE) reads per sample were sequenced on an Illumina Nextseq 500 platform in High Output (150 cycles) mode.

#### Nuclear genome analysis

Raw FASTQ reads were trimmed and filtered to remove low-quality reads (minimum base PHRED quality of 18 and minimum read length of 50bp) prior to mapping using *cutadapt* (version 2.4; (*21*)). Reads were mapped to the reference genome of *D. melanogaster* (Flybase Release 6.28) with *bwa aln* (version 0.7.12; (*22*)) using parameters optimized for Pool-seq data (*23*). Mapped reads were filtered for proper pairing and a mapping quality of at least 20 using *samtools* (version 1.9; (*24*)). Duplicates were removed with *Picard* (version 2.18.11; http://broadinstitute.github.io/picard/) and sequences flanking indels were re-aligned with *GATK* (version 3.8.1.0; (*25*)). Sequencing depth was assessed using *Qualimap* (version 2.2.1; (*26*)) and ranged from 46-58x for autosomes and 22-28x for X chromosomes. Individual *bam* files from all samples were then combined into a single *mpileup* file using *samtools* (version 1.9; (*24*)). SNPs were called with the *PoolSNP* pipeline (version 1.05; https://github.com/capoony/PoolSNP) from the DrosEU project (*27*) which was specifically developed for SNP detection in Poolseq data. Parameters for SNP calling were as those used and optimized by the DrosEU project (except minimum count was set to 10) as their dataset closely resembled the present one. The resulting *vcf* file was converted into a *sync* file using the python script *VCF2sync.py* from the DrosEU pipeline.

General genetic differentiation among nuclear genomes was assessed by principal component analysis (PCA) using the R package *LEA* (version 3.4.0; (*28*)) and by estimating admixture proportions with the R package *ConStruct* (version 1.0.4; (*29*)). Both approaches were based on major allele frequencies of nuclear SNPs on all major chromosome arms (2L, 2R, 3L, 3R, and X). In order to minimize the effects of linkage disequilibrium (LD), only SNPs at least 1kb apart and outside regions of no recombination (*30*) were considered. Major allele frequencies were calculated with the python script *sync2AF.py* from the DrosEU pipeline. Admixture proportions (K=3) for each line were inferred by non-spatial modelling with three MCMC chains per run and 10,000 iterations.

#### Mitochondrial genome analysis

Sequence data of mitochondrial genomes were retrieved from data of (*1*) who had sequenced all 27 mitonuclear lines in pools of 150 flies including a prior mitochondrial enrichment protocol. These data were reanalysed with more stringent sequencing depth criteria. *Bam* files from this previous study were combined into a single *mpileup* file using *samtools* (version 1.9; (*24*)), which was then converted into a *sync* file with *Popoolation2* (*31*). As for nuclear SNPs, a PCA using the R package *LEA* (version 3.4.0; (*28*)) was performed and admixture proportions with the R package *ConStruct* (version 1.0.4; (*29*)) were estimated based on the major allele frequencies of all mitochondrial SNPs. Major allele frequencies were calculated with the python script *sync2AF.py* from the DrosEU pipeline. Admixture proportions (K=3) for each line were inferred by non-spatial modelling with three MCMC chains per run and 10,000 iterations. Genetic differentiation for each mitochondrial SNP was assessed by estimating *F_ST_* according to (*32*) and Fisher’s exact tests to estimate the significance of the allele frequency differences. Pairwise *F_ST_* and Fisher’s exact tests per SNP were calculated between mitonuclear genotypes (replicates were pooled) with *Popoolation2* (*31*). Resulting *P*-values from the Fisher’s exact tests were corrected for multiple testing using FDR correction (*33*) and SNPs significant at an FDR<0.001 were considered as significantly differentiated between mitonuclear genotypes.

#### Quantitative trait analysis

Phenotype data were analysed in R 3.6.1. Fit of fecundity data to a negative binomial distribution was determined with firdistrplus::descdist and firdistrplus::fitdist. Generalised linear mixed models of the form

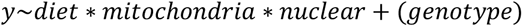

were fit with lme4::glmer.nb (egg counts & progeny, negative binomial distribution) or lme4::glmer (fertility, binomial of progeny and egg counts); in which *diet* (control/EAA/lipid), *mitochondria* (A/B) and *nuclear* (A/B) were fixed factors, genotype was a random factor denoting fly line (e.g. *AA1, AA2, AA3, AB1, AB2, BA1*, etc). Where relevant (Figure S2), experimental replicate was also included as an additional random factor. An observation-level random effect was also included for fertility under chronic feeding to ameliorate overdispersion. Anova tests (type-3) were conducted with car::Anova, and *post-hoc* analyses were applied with the functions emmeans::joint_tests, emmeans::pairs, emmeans::emmip (*14*). Options for contrasts were set to orthogonal polynomials and sum-to-zero contrasts.

Development to adult was modelled by fitting Cox mixed effects models of the form

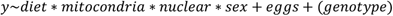

with coxme::coxme. *Diet, mitochondria, nuclear* and *genotype* terms were as in models of egg laying. *Eggs* coded number of eggs laid in the vial in which the individual developed, to account for variation in rearing density. Anova and and *post-hoc* EMM tests were conducted as per fecundity analyses.

PCA of phenotype data was conducted with prcomp on scaled EMMs, for each trait on each diet.

*r^2^* was calculated with MuMIn::r.squaredGLMM. Akaike weights were calculated with MuMIn::dredge.

To overcome challenges in calculating effect sizes for three-way DMN interactions, for each trait we used EMMs and joint tests to calculate F ratios, from which a measure of effect size within the sample population (partial η^2^) can be estimated (*14, 34, 35*). Effect sizes were calculated with custom functions built around effectsize::F_to_eta2. Application of this function was necessarily specific to the type of model in question. In all cases, F statistics were taken from ANOVA tables returned by emmeans::joint_tests. For GLMMs and Cox mixed-effect models (Figure 4), degrees of freedom were taken from ANOVA tables returned by emmeans::joint_tests; with residual degrees of freedom calculated by df.residual for GLMMs, or taken from model output for Cox mixed-effect models. For GLMs (Figure 4), degrees of freedom and residual degrees of freedom were taken from stats::anova.

EMMs superimposed on plots were calculated by fitting a GLMs of the form

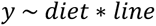

and calculating EMM per fly line per diet. A distinct model was used for EMM calculation because models used to calculate statistical effects did not return main effects for each line (e.g. coefficient for mitotype A, nucleotype A, control diet) and therefore did not show among-mitonucleotype replication (i.e. coefficients for each of lines *AA_1_, AA_2_, AA_3_* on control diet). Confidence interval of development time EMM for line *AA_3_* on +EAA diet was too large to plot owing to lethality in that condition, therefore this line was excluded from the plot (see Figure 3 legend). Values were returned to original scale by exponentiation when emmeans returned logged values.

Difference indices were calculated from EMMs per line per diet described above. For each pairwise comparison, fold-change was calculated as

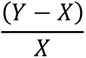

Where Y represented post-treatment value and X represented starting value. Absolutes of these values were then logged, re-signed, and scaled to a -1:1 scale. P-values for difference in EMM was calculated for each pairwise comparison using emmeans::pairs, from which FDR was returned with stats::p.adjust. Bubble plots were produced using ggplot2.

Figures were assembled in Adobe Illustrator. Mitochondria graphics were recoloured from files freely distributed under an open commons license. Skull and crossbone graphics were sourced from all-free-download.com. Heatmap of nutrient content was plotted in R with superheat.

